# Massively-parallel Microbial mRNA Sequencing (M3-Seq) reveals heterogeneous behaviors in bacteria at single-cell resolution

**DOI:** 10.1101/2022.09.21.508688

**Authors:** Bruce Wang, Aaron E. Lin, Jiayi Yuan, Matthias D. Koch, Britt Adamson, Ned S. Wingreen, Zemer Gitai

**Author notes:** Corresponding authors (B.A.); (N.S.W.); (Z.G.).

## Abstract

Bacterial populations are highly adaptive. They can respond to stress and survive in shifting environments. How the behaviors of individual bacteria vary during stress, however, is poorly understood. To identify and characterize rare bacterial subpopulations, technologies for single-cell transcriptional profiling have been developed. Existing approaches, though, are all limited in some technical capacity (e.g., number of cells or transcripts that can be profiled). Due in part to these limitations, few conditions have yet been studied with these tools. Here, we develop <u>M</u>assively-parallel <u>M</u>icrobial <u>m</u>RNA sequencing (M3-Seq), a single-cell RNA-sequencing platform for bacteria that pairs combinatorial cell indexing with *post hoc* rRNA depletion. We show that M3-Seq can profile hundreds of thousands of bacterial cells from different species under a range of conditions in single experiments. We then apply M3-Seq to reveal rare populations, insights into bet hedging strategies during stress responses, and host responses to phage infection.

## Introduction

Bacteria have a remarkable ability to survive and adapt in diverse and changing environments. One strategy that allows populations to flourish in the face of unpredictable environmental stressors is specialization of individual cells. These specializations can manifest as morphological changes (e.g., sporulation in Gram-positive organisms) or visually indistinguishable but functionally distinct states (e.g., rare antibiotic-resistant “persister” phenotypes in *Staphylococcus aureus* and *Escherichia coli*)^1–3^. A promising approach to study such specializations is to measure how single cells orchestrate gene expression in natural growth settings and in response to perturbations. For mammalian cells, such efforts have been greatly enabled by single-cell RNA sequencing (scRNA-seq)^4–6^; however, despite pioneering efforts to develop similar tools for bacteria, current technologies lag behind.

Existing bacterial scRNA-seq methods include MATQ-Seq^7^, PETRI-Seq^8^, microSPLiT^9^, par-SeqFISH^10^ and a droplet-based method^11^ (Fig. 1A and Table S1). Each of these methods uses different strategies to index cells and their transcripts, and each has associated benefits and drawbacks^12^. MATQ-Seq isolates single cells into separate wells of multiwell plates and performs individual reverse transcription and indexing reactions to generate sequencing libraries^7^. This ‘indexing’ scheme, although straightforward, is inherently limited in scale. By contrast, each of the remaining methods allows single-cell gene expression to be profiled across pools of cells in single experiments, with multiplexed transcript detection enabled by *in situ* probe hybridization (SeqFISH and the method pioneered by McNulty and colleagues) or split-pool combinatorial indexing^6^ (PETRI-Seq and microSPLiT). However, drawbacks remain. Hybridization-based approaches rely on pre-designed species- and gene-specific probes which limit unbiased discovery, while the combinatorial indexing platforms suffer from an abundance of signal from ribosomal RNA (rRNA), which can compromise mRNA detection. Given these considerations, we developed <u>m</u>assively-parallel <u>m</u>icrobial <u>m</u>RNA-<u>seq</u>uencing from single cells (M3-Seq), a method for scRNA-seq in bacteria that combines plate-based, *in situ* indexing with droplet-based indexing and *post hoc* rRNA depletion. M3-Seq enables gene expression profiling of hundreds of thousands of single bacteria across many samples at transcriptome-scale with sensitive mRNA capture. By applying M3-Seq, we reveal independent phage induction programs in *Bacillus subtilis (B. subtilis*), a bet-hedging subpopulation of *Escherichia coli* (*E. coli*), heterogeneities in multiple species, and hostpathogen interactions after phage infection.

**Figure 1.**
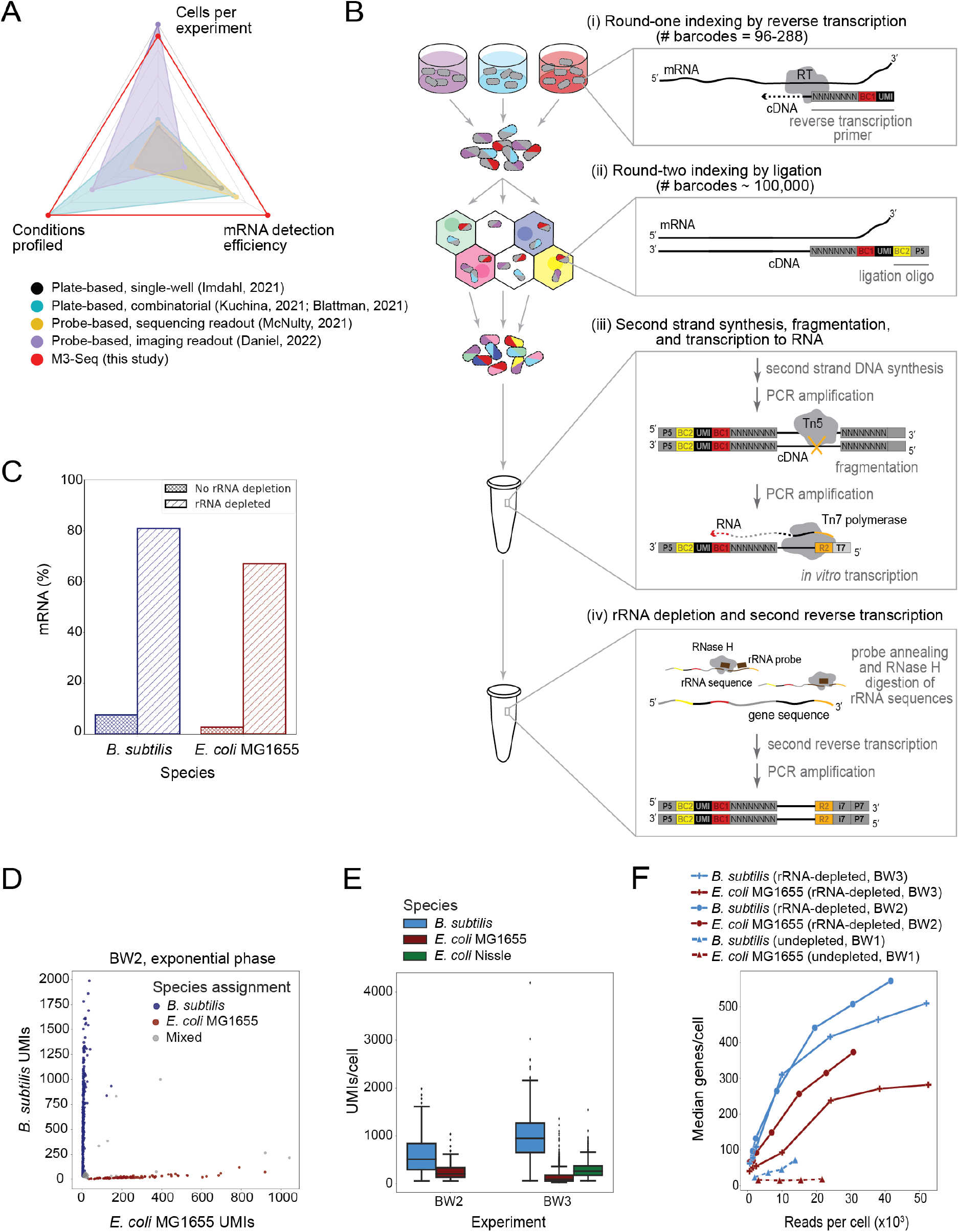
Development of M3-Seq platform for single-cell RNA-sequencing with *post hoc* rRNA depletion. **A.** scRNA-seq methods previously established for bacteria with reported number of cells (ranging from 100 cells/experiment – 300,000 cells/experiment), conditions (ranging from 1 condition/experiment to 20 conditions/experiment), and mRNA genes per cell (ranging from 29 genes/cell to 371 genes/cell). Numbers in each category were selected by taking maximum reported values. Numbers also found in Table S1. **B.** Schematic of M3-Seq experimental workflow. Indexing: (i) RNA molecules are reverse transcribed *in situ* with indexed primers such that cells in each reaction (i.e., separate plate wells) are marked with distinct sequences. Primers allow for random priming. (ii) Cells are then collected, mixed, and distributed into droplets for a second round of indexing via ligation with barcoded oligos. Sequencing library preparation: Cells are collected again and lysed to release single-strand cDNAs. (iii) Second strand synthesis is then performed in bulk reactions and resulting cDNA molecules are fragmented with Tn5 transposase, amplified via PCR to add a T7 promoter, and converted to RNA using T7 RNA polymerase. (iv) To deplete ribosomal sequences, the amplified RNA library is hybridized to complementary DNA probes (Table S4), and annealed sequences are cleaved by RNase H. Finally, remaining sequences are reverse transcribed back to DNA, sequencing adaptors are added, and data is collected by sequencing. **C.** Percentages of mRNA sequences in *B. subtilis* and *E. coli* single-cell libraries prepared with and without rRNA depletion (>20-fold more mRNA observed with depletion). Data from undepleted libraries come from BW1, and data from depleted libraries come from BW3. **D.** M3-Seq analysis of a mixture of *B. subtilis* and *E. coli* wherein each point corresponds to a single “cell” (i.e., unique combination of plate and droplet barcodes). Species assignments were made if >85% of UMIs mapped to unique species-specific transcripts. Otherwise, cells were designated as mixed. **E.** UMIs per cell (after species assignment) observed in exponential phase cells across two experiments, BW2 and BW3 (515 ± 245 and 953± 310 median UMIs with absolute deviation for *B. subtilis*, respectively; 211 ± 85 and 100±47 median UMIs with absolute deviation for *E. coli* MG1655, respectively; 266±100 UMIs with for *E. coli* Nissle in BW3). **F.** Median genes detected per *B. subtilis* or *E. coli* cell as a function of number of total reads per cell across three experiments, BW1, BW2, and BW3. rRNA depletion in two experiments (solid curves, BW2, BW3) enabled an order of magnitude greater detection than without that step (dashed curves, BW1).

## Results

### M3-Seq captures rRNA-depleted transcriptomes from single bacterial cells

We designed M3-Seq to build on scifi-RNA-seq^13^, a combinatorial indexing platform for mammalian cells. M3-seq has two rounds of cell indexing (Fig. 1B and S1). The first of these indexing rounds uses *in situ* reverse transcription with random priming to tag transcript sequences with one cell index (BC1) and a unique molecular identifier (UMI). This indexing step, which we refer to as “round-one indexing”, occurs in multiple reactions, each performed on a separate pool of fixed, permeabilized bacterial cells. After this step, cells are mixed and then separated again into droplets using a commercially available kit (Chromium Next GEM Single Cell ATAC, 10X Genomics). In these droplets, we perform “round-two indexing”, wherein a second cell index (BC2) is ligated onto cell-associated, BC1-indexed cDNA molecules. While neither BC1 nor BC2 are necessarily unique, together these sequences create a combinatorial index that should serve as a distinct marker for individual cells, even when multiple cells are indexed within a single droplet during round-two. Conceptually, this indexing scheme is identical to scifi-RNA-seq^13^, which enabled sequencing of >100,000 mammalian cells in a single run. However, because bacteria are considerably different than mammalian cells (e.g., smaller, thick cell walls), we performed a series of pilot experiments before testing the scheme at scale. First, we verified that we could load singlecell suspensions of bacterial cells into droplets at rates appropriate for combinatorial indexing. Specifically, we loaded different numbers of Sytox Green-stained *E. coli* into droplets, calculated cell loading distributions by imaging (Fig. S2A, B), and determined expected index collision rates across numbers of round-one indices (Fig. S2C). These calculations suggested that with ~96 round-one indices, hundreds of thousands of cells can be loaded in a single run of the droplet system with <1% collision rate between combinatorial BC1 and BC2 indices.

Next, we verified that, even though bacterial cells are surrounded by thick cell walls and contain very few mRNA molecules, we could generate single-cell transcriptomes using our approach. Briefly, after growing *B. subtilis* 168 and *E. coli* MG1655 to exponential and stationary phase, we fixed, washed, and permeabilized the cells with lysozyme^8,9^. We pooled the cells at equal cell numbers, performed combinatorial indexing using 96 round-one indices, and loaded 100,000 cells into droplets for round-two indexing (1 lane of a 10x genomics run). We refer to this experiment as BW1 (Table S2 and S3). Given our previous loading calculations, we would expect 15.7% of all cell-containing droplets in this experiment to yield an index collision without round-one indexing and, similar to expectations, our data revealed a 12.7% collision rate between *B. subtilis* and *E. coli* cells when only BC2 indices were used to discriminate cells (25.6% when accounting for within-species collisions^14^). Encouragingly, using both BC1 and BC2 indices dramatically decreased this rate to 0.7% (1.5% when accounting for within-species collisions) (Fig. S2D, E). Moreover, demonstrating that our approach can generate single-cell transcriptomes, comparison of average profiles from exponential phase *E. coli* were similar to published bulk RNA-seq data^8^ (*r* = 0.59) (Fig S2F).

As has been previously observed with other bacterial combinatorial indexing methods^8,9^, most reads in our pilot experiment aligned to rRNAs (Fig. S2G-J). For example, of roughly 1,000-2,000 reads per cell in exponential phase *E. coli*, only ~100 (3-10%) aligned to mRNA, and the rest aligned to rRNA (90-97%). While this problem can be overcome by sequencing to greater depth, we sought a more cost-effective solution and thus developed steps to remove rRNA sequences prior to sequencing. When developing these steps, we considered the observation that depletion of rRNA *in situ* can decrease mRNA capture efficiency^9^ and thus focused on depleting rRNAs after amplification (Fig. 1B and S1). Specifically, after testing two approaches for depleting ribosomal sequences from bulk libraries (Fig. S3A, B), we chose an RNase H-based approach^15–17^ to complete our pipeline (Fig. 1A and S1A). Altogether our full M3-Seq pipeline is as follows: After two rounds of indexing (performed as described above), cDNA libraries are transcribed to singlestranded RNA. rRNA sequences within the library are then hybridized to rRNA-specific DNA probes and digested with RNase H, which specifically cleaves RNA in RNA:DNA hybrids (Fig. S1B). Resulting rRNA-depleted libraries are then reverse transcribed back into cDNA for sequencing. Encouragingly, putting these steps together enabled recovery of single-cell transcriptomes with 66.9% of *E. coli* reads and 80.8% of *B. subtilis* reads aligning to mRNA, a >20-fold increase from our previous experiment (Fig. 1C). Furthermore, we observed that the mRNA content of our rRNA-depleted bulk libraries was similar to libraries that has not been depleted (*r* = 0.77) (Fig. S3B), implying that biological signal is not lost during depletion.

To evaluate the full M3-Seq pipeline, we next performed two large experiments (Table S2 and S3): one in which we evaluated *B. subtilis* 168 and *E. coli* MG1655 (BW2) and one in which we evaluated these species alongside the probiotic *E. coli* strain Nissle 1917 (BW3). In both of these experiments, we grew bacteria to exponential (OD=0.3) and early stationary phases (OD=2.5, 2.8, and 2.6) with and without antibiotic treatments. After in-plate, round-one indexing, we pooled cells from each condition and loaded them into droplets. From BW2, we recovered 4,975 singlecell transcriptomes from 37,500 loaded cells (13% recovery rate), and from BW3, we recovered 15,539 single-cell transcriptomes from 67,670 loaded cells (22% recovery rate) (Table S3). Consistent with our previous experiments, we observed a low index collision rate among samples loaded into droplets in the same reaction (Fig. 1D, S3C-E). After identifying single cells using combined round-one and round-two indices, discriminating samples by round-one indices, and distinct species using the aligned mRNA transcripts, we recovered 515 and 984 median UMIs per exponential phase *B. subtilis* cell (298 and 371 median genes per cell), 211 and 100 median UMIs per exponential phase *E. coli* MG1655 cell (75 and 151 median genes per cell), and 266 median UMIs per exponential phase Nissle cell (175 median genes per cell), respectively (Fig. 1E and S3F). Compared to other studies that applied scRNA-seq to bacteria^8,9,11^, this represents roughly the same number of UMIs per cell for *E. coli* but twice as many UMIs per cell for *B. subtilis*. Critically, data from these experiments also revealed that M3-Seq libraries require ~15X fewer reads per cell to detect the same number of genes as un-depleted libraries (Fig. 1F). Moreover, similarity between one *E. coli* sample and an existing RNA-seq dataset^8^ (*r* = 0.72) (Fig. S3G) indicated that the bulk biological signal is retained throughout our experimental pipeline, and we found that biological replicates had similar subpopulation compositions (Fig. S3H-J) and that bulk biological signal was retained between replicates (*r* = 0.94, 0.79, 0.92) (Fig S3K-M). Thus, by combining our *post hoc* rRNA depletion with droplet overloading, M3-Seq provides biologically meaningful, mRNA-enriched transcriptomes at single-cell resolution.

### Early stationary phase *E. coli* contain a rare, acid-tolerant subpopulation

The transition from exponential phase to early stationary phase represents a shift from rapid cell division to slow growth as nutrients are depleted from the environment. Across the three bacterial strains in our BW3 experiment, our single-cell data successfully distinguished stationary phase cells from those that were growing exponentially, as demonstrated by unbiased separation of cells marked by growth-stage-specific “round-one” indices (Fig. S4A-C) and gene ontology (GO) analysis of genes differentially expressed between those cells, which showed clear enrichment for biological processes associated with one growth stage or the other (Fig. S4D-F). Additionally, as would be expected from dampened transcriptional output during slowed growth, stationary phase cells had substantially fewer UMIs per cell than exponential phase samples, with a median of 30 UMIs per cell for *B. subtilis* and *E. coli* MG1655, and 39 UMIs per cell for Nissle.

Strikingly, in addition to differences between cells collected at different growth stages, we observed transcriptional heterogeneity *within* populations of early stationary phase cells (Fig. 2A-B, Fig. S5A-B). A closer examination of cells from both strains of *E. coli* in this growth stage revealed clusters of cells overexpressing genes involved in intracellular pH elevation and glutamate catabolism (Fig. 2C, Fig. S5C). The most strongly expressed genes in these clusters were *gadA* and *gadB* (Fig. 2D-E, Fig S5D-E). These genes are well conserved among enteric bacteria and are known to encode glutamate decarboxylases that de-acidify the cellular cytoplasm by consuming a proton during decarboxylation of glutamate to GABA (Fig. S5F)^18–20^. Notably, previous studies have shown that these genes are expressed in stationary-phase *E. coli* using bulk measurements^21,22^, and heterogeneous expression has been observed in other conditions^23,24^. However, heterogeneous expression of *gadA* and *gadB* during the transition into stationary phase has not been previously reported. To confirm this finding, we transformed *E. coli* MG1655 with a plasmid encoding a fluorescent reporter controlled by the *gadB* promoter (P_*gadB*_-GFP) and imaged the cells after growth in the same condition used for single-cell sequencing (Fig. 2F, inset). Analysis of the resulting images revealed a heavy tail of GFP expression in the population, indicative of a small subpopulation of cells with active *gadB* transcription (Fig. 2F).

**Figure 2.**
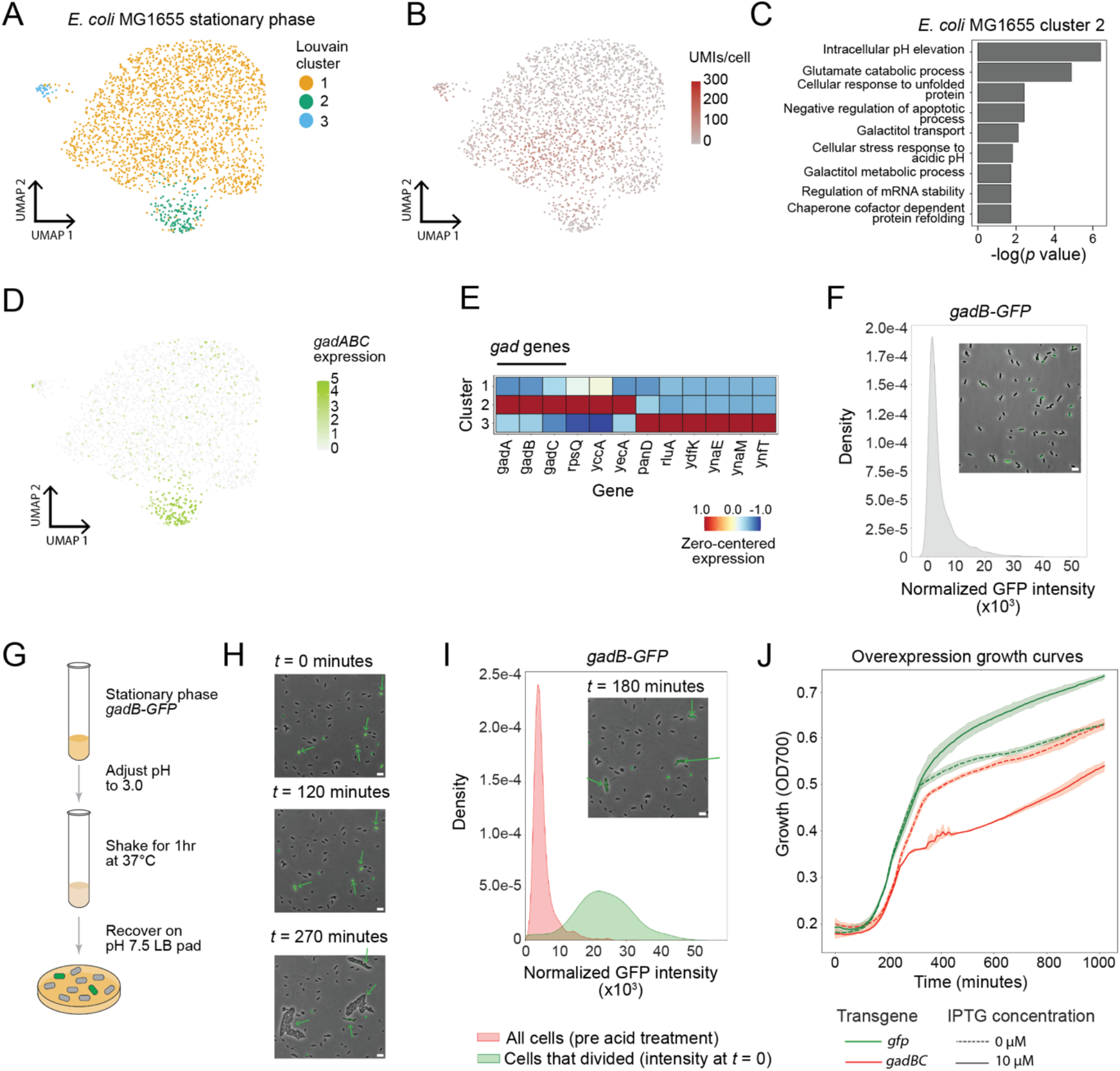
M3-Seq reveals an acid-tolerant, bet-hedging subpopulation of *E. coli* in early stationary phase. **A.** UMAP projection of *E. coli* MG1655 transcriptomes from cells at early stationary phase (OD=2.8). Colors indicate clusters of transcriptionally similar cells. **B.** Same as (A) but with color gradient indicating number of UMIs captured in each cell. **C.** GO-term enrichment of select biological processes calculated with marker genes identified for cluster 2 in (A). Marker genes were determined by comparing the within-cluster average expression to out of cluster average expression and filtering for genes with *p*-value < 0.05 (Wilcoxon-rank sum test). The *p*-values are -log_10_ transformed such that the most strongly enriched biological processes have the highest score. Selected processes were those with the lowest *p*-values (Fisher’s exact test) after thresholding at 0.05. **D.** Same as (A) but with color gradient indicating expression of *gadABC* genes. **E.** Zero-centered and normalized expression of marker genes for each cluster identified in (A). Marker genes were defined as those observed in at least 5% of cells in that cluster, and with the lowest *p*-values (Wilcoxon rank-sum test) after thresholding to select genes with >0.5 log_2_ fold change between within-cluster and out-of-cluster average expression. A maximum of 6 gene were included per cluster. **F.** Normalized fluorescence distribution of *E. coli* transformed with P_*gadB*_-GFP grown in LB media to OD=2.8. Inset is a representative composite image with phase and GFP channels overlaid. Scale bar, 5 *μ*m. **G**. Schematic of acid-stress recovery assay. Cultures grown as in (F) were adjusted to pH = 3.0 using 12N HCl, allowed to shake for 1 hour, and placed onto a fresh LB-agarose pad at pH 7.5 for imaging over 9 hours. **H.** Representative composite images of *E. coli* expressing P_*gadB*_-GFP during recovery phase of acid-stress recovery assay depicted in (G). t=0 represents time of placement onto fresh LB-agarose pad. Arrows indicate cells that divided during the recovery period. Scale bar, 5 *μ*m. **I.** Kernel density estimates of the fluorescence distribution of *E. coli* expressing P_*gadB*_-GFP before and after acid-stress recovery assay. Red depicts measurements from cells before acid treatment. Green depicts measurements from cells at *t* = 0 that ultimately divided over the course of 8 hours of recovery (i.e., survived acid treatment). Inset is a representative composite overlay of the cells 180 minutes after the start of recovery from the same experiment as in (H). Arrows indicate cells that divided during the recovery period. Scale bar, 5 *μ*m. **J.** Growth curves of *E. coli* MG1655 transformed with GFP or *gadBC* transgene (overexpression plasmids) and grown with or without 10 *μ*M of IPTG (dashed lines) for 1000 minutes. Induction of *gadBC* (red) reduced growth compared to the uninduced, whereas induction of GFP (green) did not.

Our finding that *gad* genes are heterogeneously expressed in early stationary phase presented an opportunity to experimentally validate our single-cell data and, at the same time, investigate the function of heterogeneous gene expression during a biologically important process. To confirm a functional role for the *gad* genes in our cells, we asked if *E. coli* MG1655 lacking *gadABC* can survive acid stress applied during early stationary phase. Data from this experiment, which measured the number of viable cells by counting colony forming units (CFU) with and without acid stress revealed that acid tolerance in the triple deletion strain was strongly impaired relative to wildtype (Fig. S5G). However, given the experimental design, these data could not link surviving cells to any pre-existing subpopulation. We therefore next deployed our P_*gadB*_-GFP reporter strain to monitor how cells with varying levels of *gadB* expression recover from acid treatment (Fig. 2G-H). First, we grew the reporter strain to early stationary phase and using imaging, confirmed that a subpopulation of the cells expressed the *gad* genes. Next, we exposed the whole population of cells to acid stress (pH 3.0) and after 1 hour, transferred an aliquot of the stressed cells to a fresh LB agarose pad. We then imaged the cells for 8 hours, and quantified GFP intensity as a proxy for *gadB* expression across individual cells in pre- and post-treatment images (Fig 2I). Analysis of these quantifications revealed an intriguing finding: the population of viable cells, which we define as cells that divided at least once during the recovery period, were those already expressing high levels of GFP at the beginning of the recovery. This observation hints that the subpopulation of cells expressing *gadB* before acid exposure are the ones that subsequently survived acid treatment. To further explore this possibility, we imaged early stationary P_*gadB*_-GFP reporter cells during strong acid stress and found that, rather than increasing in response to treatment, GFP fluorescence intensity steadily decreased in bacterial cells (Fig. S5H, Movie 2). Together, these observations suggest that under sudden strong acid stress, *E. coli* do not activate and translate acid resistance proteins, but instead rely on an existing pool of translated proteins in a subpopulation of cells.

A reason for having only a subpopulation of cells express the *gad* genes at levels that are protective against acid stress would be if there is a significant cost to expressing these genes. To test this hypothesis, we cloned the *gadBC* operon into an inducible-overexpression vector and performed growth-curve assays in LB across a range of inducer concentrations^25,26^. This experiment revealed that overexpression of *gadBC* at even low induction levels (1-10 μM, which is 50-1000x less than typical induction concentrations) causes a growth defect (Fig. 2J, Fig. S5I). Paired with our functional characterization of the *gadB*-expressing subpopulation, these data suggest a model wherein *E. coli* can preemptively activate the *gad* genes to protect against future strong acid stresses (e.g., such as would be experienced when passing through acidic environments like the stomach), but because *gad* expression causes decreased growth overall, activation can be limited to a subpopulation in case the acid stress does not materialize. We therefore conclude that during entry to stationary phase, enteric bacteria asynchronously activate the *gad* genes as a bet-hedging strategy to protect some cells against strong acid stress while enabling other cells to grow rapidly.

### Identification of multiple transcriptional states in *E. coli* treated with ribosome-targeting, bacteriostatic antibiotics

How bacteria respond to antibiotic-treatment is an important question. However, the large number of bacterial species and types of antibiotics, combined with variability of response within populations, makes this a difficult question to approach systematically. Combinatorial indexing provides a straightforward way to evaluate gene expression across many samples (i.e., separate round-one indices can mark many cultures) and given the single-cell resolution of our platform, we reasoned that M3-Seq could prove beneficial in this space. We therefore deployed M3-Seq to evaluate bacterial cultures treated with each of eight antibiotics: two DNA damaging agents (nalidixic acid, ciprofloxacin), two inhibitors of cell wall synthesis (cycloserine, cefazolin), and four ribosomal inhibitors (chloramphenicol, erythromycin, tetracycline, gentamycin) (Fig. 3A, Table S2 and S3). In this experiment (BW4), cultures were grown to early exponential phase (OD=0.3), treated with 2X the minimum inhibitory concentration of each drug for 90 minutes, and subjected to M3-Seq across 2 lanes of a 10X Genomics scATAC run. Altogether, we report data for 20 conditions across 229,671 cells (Table S3), from which we make two systems-level observations: First, indicative of successful profiling, select genes with known associations to antibiotic-induced stresses had higher expression in expected cultures (Fig. S6A, B). Second, hierarchical clustering of correlations between pseudobulk expression profiles grouped drugs with the same mechanism of action, suggesting that M3-Seq is a promising tool for systematic analysis (Fig. 3B, C).

**Figure 3.**
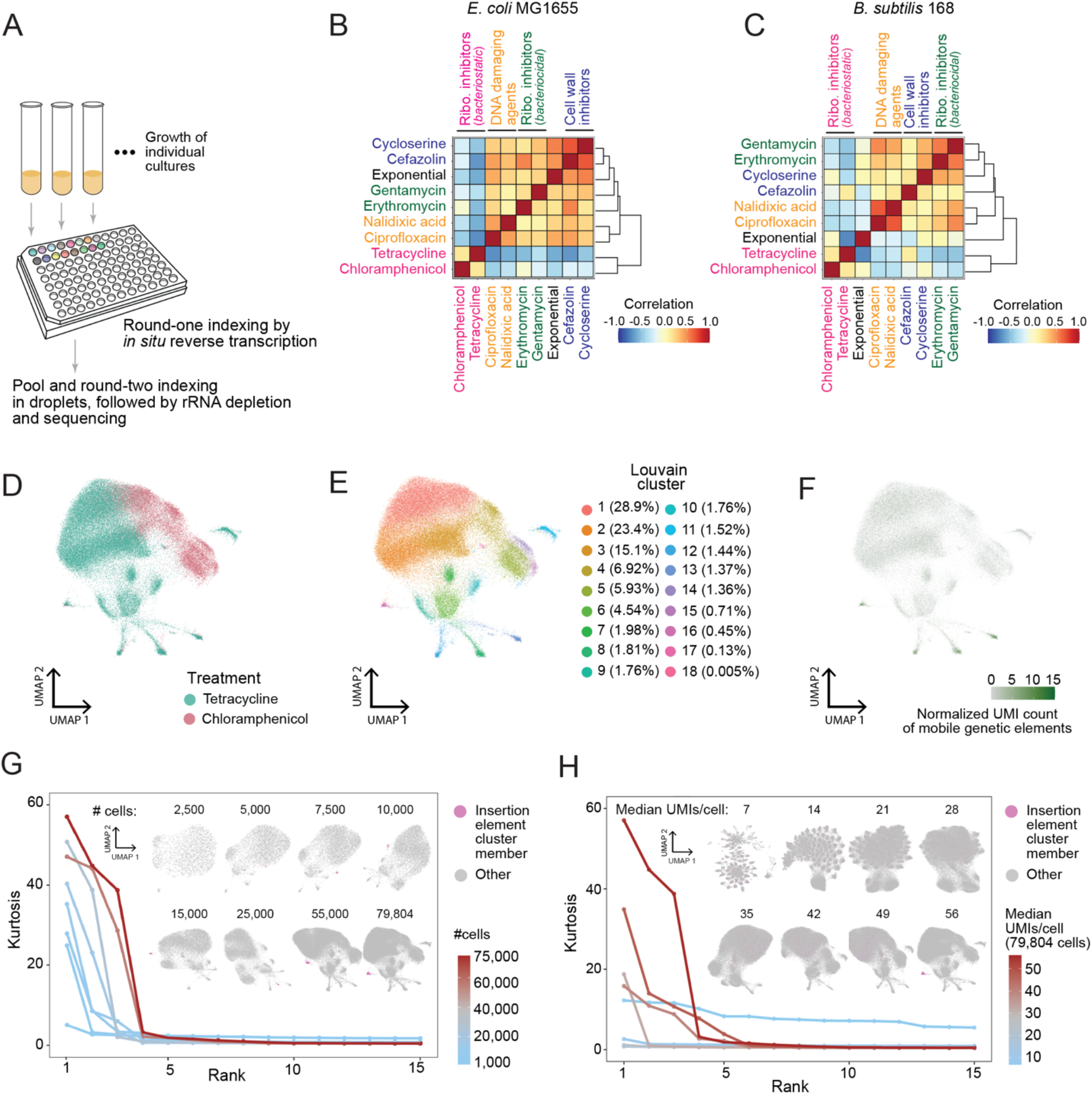
M3-Seq enables systematic investigation of bacterial response to antibiotic treatment. **A.** Schematics of antibiotic experiment (BW4). During preparation of M3-Seq gene expression libraries, round-one plate indexing was used to uniquely mark antibiotic-treated and untreated cultures. **B.** Heatmap depicts Pearson correlations of pseudobulk transcriptomes from *E. coli* MG1655 prepared as in (A), which were computed using genes with average expression greater than the median average expression of all genes. Colored text indicates antibiotics of similar mechanisms of action. **C.** Same as (B) but for *B. subtilis* 168. **D.** UMAP projection of *E. coli* MG1655 transcriptomes from cells treated with the bacteriostatic antibiotics tetracycline and chloramphenicol. Colors indicate drug treatment. **E.** Same as (D) but with colors indicating clusters of transcriptionally similar cells. Percentage of cells in each cluster denoted. **F.** Same as (D) but with color gradient indicating the normalized UMI count of mobile genetic elements. Clusters 8, 12, 13,16 were enriched for MGE expression. **G.** Cell rarefaction analyses using M3-Seq data. Line graph indicates kurtosis of 15 principal components computed from tetracycline- and chloramphenicol-treated *E. coli* MG1655 cells, with individual lines corresponding to calculations from the total population of cells (79,804) or down-sampled populations thereof (down to 1,000 cells). The 15 principal components included were those with the highest kurtosis. Inset UMAP projections were computed from each down-sampled data matrix. Within the embeddings, magenta indicates members of cluster 16 (indicated in F), which can only be distinguished >7,500 – 10,000 cells. Notably, the top row of embeddings (2500 – 10,000 cells) represents the scale of experiments from previous studies, while the bottom row represents the scale enabled by M3-Seq. **H.** UMI rarefaction experiments using M3-Seq data. Line graph indicates kurtosis of 15 principal components computed from 79,804 tetracycline- and chloramphenicol-treated *E. coli* MG1655 cells, with individual lines corresponding to data subsampled for UMIs per cell (7 to 56 median UMIs). The 15 principal components included were those with the highest kurtosis. Inset UMAP projections were computed from each downsampled data matrix. Within the embeddings, magenta indicates members of cluster 16 (indicated in F), which can only be distinguished at the highest UMI detection efficiency.

A closer examination of individual samples at the single-cell level (Fig. S6C, D) next revealed that tetracycline- and chloramphenicol-treated *E. coli* had a large number of transcriptional states (14 and 8 clusters, respectively) (Fig. S6C, Table S5). Unlike bactericidal drugs, such bacteriostatic agents do not have readily-measurable single-cell persistence and tolerance phenotypes^2,27–29^, and thus relatively little is known about heterogeneity in response to these drugs. Exploring the data from these conditions together, we then identified several rare clusters of cells which contained cells from both samples and expressed genes from mobile genetic elements (MGEs) (Fig. 3D-F, S7A-D, Table S5). Such rare cell populations may help cultures tolerate and escape the bacteriostatic state through subtle mechanisms (e.g., increasing genetic diversity via the insertion element *insI-2* or activating genes implicated in cold shock, such as *ydfK*). From a technical perspective, though, these samples also provided the largest number of transcriptomes and among the highest median UMIs per cell from our experiment (Fig. S6C). We would therefore expect higher sampling of rare cell populations and better detection of genes from those conditions. The large number of cells (79,804 from the two conditions combined) and high median UMIs (55 and 65 for tetracycline- and chloramphenicol-treated samples, respectively) within these populations thus also provided an opportunity to evaluate requirements of scale and mRNA capture.

To better understand how the ability to detect rare subpopulations increases with the number of cells sequenced and UMIs captured, we first needed a metric capable of capturing transcriptional variability in the data. We found in our data that certain principal components had heavy tails and that cells in these tails were assigned as members of unique subpopulations in our clustering analysis (Fig. S7E-H). We therefore reasoned that we could assess detection of rare cell subpopulations by computing the kurtosis (a measure of how heavy the tails of a distribution are) for each principal component (Fig. S7I-J)^30^. Performing this analysis on down-sampled versions of the data showed that the kurtosis of the top principal components (ranked by kurtosis) decreased when the data was down-sampled (Fig. 3G-H). Correspondingly, a cluster containing the rare cell populations expressing *insI-2*, was undetectable when the data were down sampled, with no ability to detect at lower cell numbers and UMI capture rates, including those relevant to other samples from this experiment, as well as previous studies (~1,000 – 5,000 cells, 7-49 UMIs/cell). This population nevertheless became apparent above our down sampling of 7,500 cells and 56 UMIs/cell. Notably, the kurtosis of the “heaviest” tailed principal components monotonically increased with increasing cell numbers up to the number of cells in our experiment (79,804 cells) and the number of median mRNA transcripts captured (56 UMIs), suggesting that sequencing even more cells with deeper mRNA coverage could potentially identify even rarer subpopulations. Our combined analysis thus illustrates the need for scRNA-seq analysis to be performed at massive scale in bacteria and show how M3-Seq can enable such efforts.

### DNA damaging antibiotics independently induce two distinct prophages in *B. subtilis*

A second observation from our antibiotic study was that in cells treated with DNA damaging agents, *B. subtilis* exhibited heterogeneous expression of genes associated with either of two prophages present within the *B. subtilis* genome (Fig. 4A-F, Fig. S6D). While these prophages (PBSX and SPβ) are well known to be induced by conditions that induce the SOS response (such as DNA damage^31^), our single-cell data provided the opportunity to address an outstanding question: At the level of individual cells, is prophage induction stochastic or determined by some common perturbation (i.e., degree of damage) or cross-talk (i.e., co-repression)? Suggestive of the former, clustering analysis separated prophage-expressing cells into three groups: one dominated by PBSX-expressing cells (cluster 5) and two dominated by SPβ -expressing cells (cluster 6 and 7) (Fig. 4F, Table S5). Further, on a per cell basis, comparison of PBSX and SPβ transcript percentages showed no obvious correlation (Fig. 4G) and rates of co-induction across cells, which we determined by thresholding, closely matched an assumption of independence (2.44% observed, 2.47% expected) (Fig. 4H). Therefore, we found no evidence for cross-repression or results supporting a model wherein individual cells with the greatest damage had the greatest likelihood of inducing both prophages. Validation of prophage induction using single-molecule FISH (smFISH) on ciprofloxacin-treated cells, which we performed with probes against the most strongly expressed PBSX and SPβ genes, further supported this conclusion (Fig. 4I).

**Figure 4.**
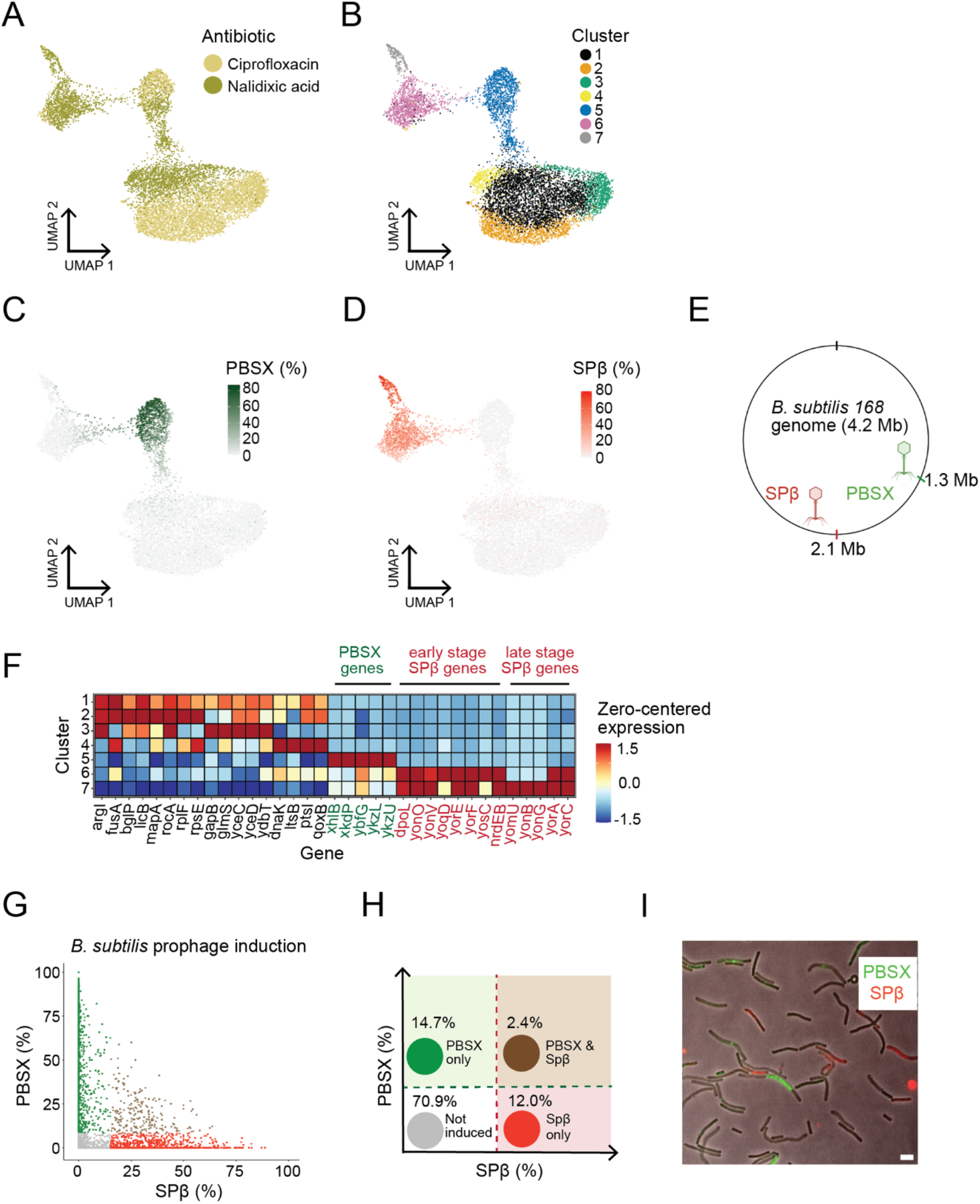
M3-Seq characterizes independent activation of prophages in *B. subtilis*. **A.** UMAP projection of *B. subtilis* transcriptomes from ciprofloxacin- and nalidixic acid-treated cells in exponential phase (OD = 0.3). Colors indicate treatment conditions (90 minutes). **B.** Same as (A) but with colors indicating clusters of transcriptionally similar cells. **C.** Same as (A) but with color gradient indicating percentage of PBSX prophage UMIs within each cell. Percentages were calculated by dividing the total number of PBSX UMIs by the total number of UMIs in each cell. **D.** Same as (A) but with color gradient indicating percentage of SPβ prophage UMIs within each cell. **E.** Schematic of *B. subtilis* genome with location of PBSX and SPβ prophages indicated. **F.** Zero-centered and normalized expression of marker genes for each of seven clusters identified in (B). Marker genes were defined as those observed in at least 5% of cells in that cluster, and with the lowest *p*-values (Wilcoxon rank-sum test) after thresholding to select genes with >0.5 log_2_ fold change between within-cluster and out-of-cluster average expression. A maximum of 5 gene were included per cluster. PBSX and SPβ prophage genes were upregulated in clusters 5, 6, 7. **G.** Left: Classification of cells with induced prophages. Green indicates cells with relative expression of PBSX genes greater than >8.4% per cell, which is >10^th^ percentile of PBSX prophage gene expression in cluster 5 from (B). Red indicates cells with relative expression of SPβ genes greater than >15.0% per cell, which is >10^th^ percentile of SPβ prophage gene expression in cluster 6 from (B). Brown indicates cells above both thresholds. Right: Schematic of prophage classification results. The expected independent coinduction probability (calculated from observed PBSX and SPβ percentages) is 2.5%. **H.** Dual color smFISH of *B. subtilis* treated with ciprofloxacin for 90 minutes. Probes hybridizing to PBSX genes were labeled with a green fluorophore. Probes hybridizing to SPβ genes were labeled with a red fluorophore. Scale bar, 5 *μ*m.

### M3-Seq enables the study of host-pathogen interactions at the single-cell level

After observing gene expression from prophages using M3-Seq, we reasoned that the platform could also be useful for studying active phage infection. Previous studies have evaluated transcriptional responses to phage with bulk measurements^32,33^. However, the cell-to-cell variability of phage adsorption and infection limits our ability to interpret these data^34–36^. To address this limitation, we characterized gene expression in individual *E. coli* cells after infection with λ phage. We conducted this experiment alongside the antibiotics experiment (BW4). Briefly, we infected exponential phase *E. coli* MG1655 (grown to OD=0.3) with λ phage at multiplicity of infection (MOI) of ~100 (Fig. S8A). We sampled the cultures at 30- and 90-minutes post infection, performed M3-Seq, and aligned the sequencing reads to a combined *E. coli* and λ genome. Cell transcriptomes formed four distinct clusters, with one cluster (3) demonstrating high levels of λ gene expression (Fig. 5A-D). During lysis, λ overtakes the host transcriptional machinery to express high levels of the late-stage genes required to produce functional virions. Indicative of lytic infection, cluster 3 revealed particularly high levels of late stage λ genes (i.e., *H, A, B, E, J, K*) (Fig. 5E, Fig. S8B, Table S5). By contrast, the most highly expressed genes in the remaining clusters (1, 2, and 4) were from *E. coli* (Fig. 5C, Table S5). Given the saturating MOI used in the experiment, our expectation was that all cells would be infected, but because cluster 3 represents a minority of the cells in the experiment (13.1% and 9.82% of the 30- and 90-minute samples, respectively), these data demonstrate how even at high MOIs, bulk measurements do not accurately reflect processes occurring during phage infection.

**Figure 5.**
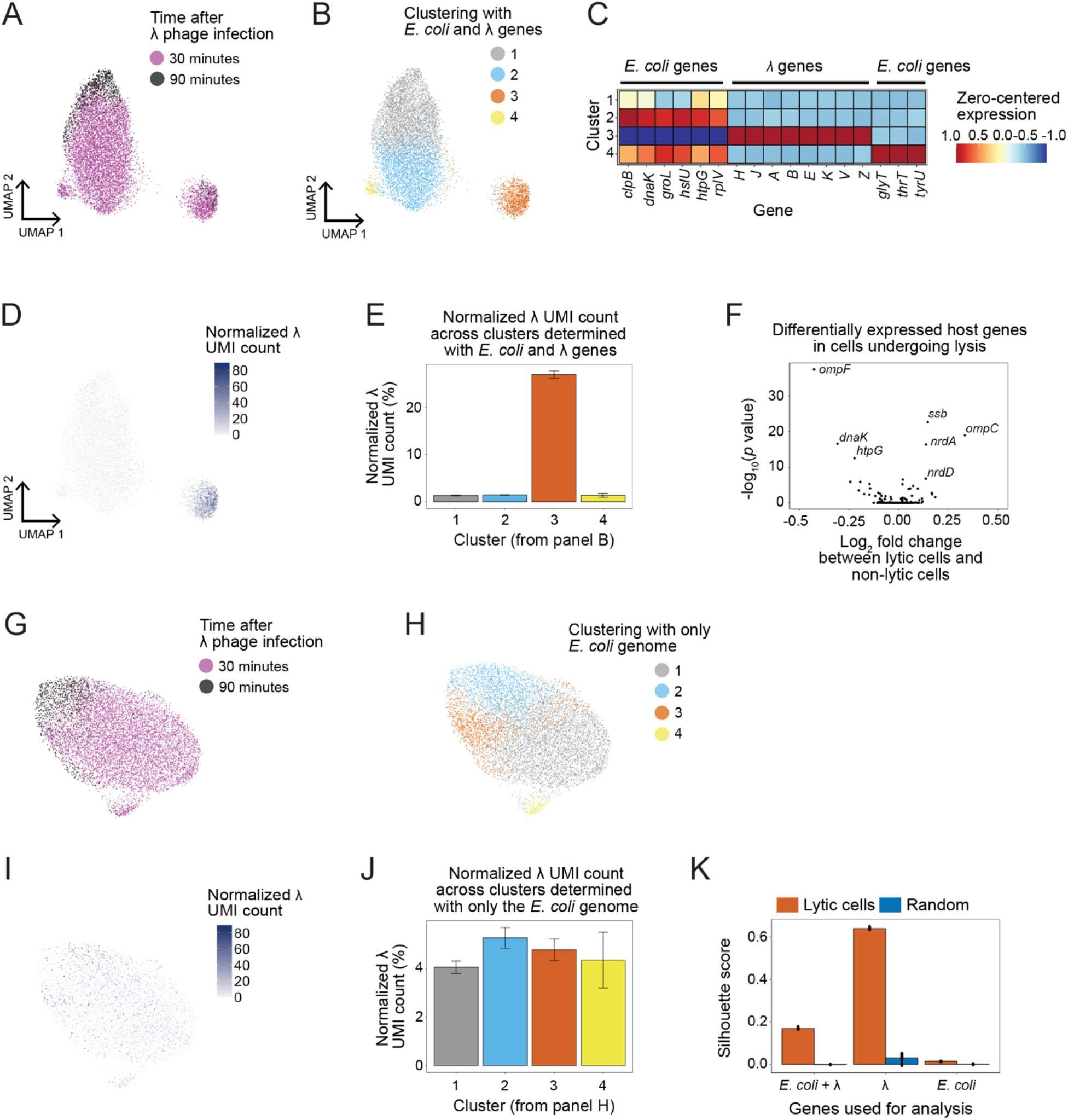
M3-Seq reveals limited host response to heterogeneous λ phage infection. **A.** UMAP projection of phage infected cells generated using alignments to both *E. coli* MG1655 and λ phage genomes (“combined genome”). Colors indicate sampling timepoint after infection. **B.** Same as (A) but with colors indicating clusters of transcriptionally similar cells. **C.** Zerocentered and normalized expression of marker genes for each of four clusters identified in (B). Marker genes were defined as those observed in at least 5% of cells in that cluster, and with the lowest *p*-values (Wilcoxon rank-sum test) after thresholding to select genes with >0.5 log_2_ fold change between within-cluster and out-of-cluster average expression. A maximum of 6 gene were included per cluster. Marker genes for cluster 3 correspond to late-stage λ lytic genes. **D.** Same as (A) but with color gradient indicating normalized λ phage UMI count in each cell. Cluster 3 from (B) is strongly enriched for λ transcripts. We refer to this group of cells as the “lytic cluster”. **E.** Normalized λ UMI count across each cluster in (B). **F.** Volcano plot of all host genes when comparing the cells in the lytic cluster to cells outside the cluster. Data show minimal log_2_ fold changes. Fold changes and *p*-values were computed using the FindMarkers function in Seurat, where the “min.pct”0 and “logfc.threshold” were both set to 0. **G.** UMAP projection of phage infected cells generated using alignments to only the *E. coli* MG1655 genome. Colors indicate sampling timepoint after infection. **H.** Same as (G) but with colors indicating clusters of transcriptionally similar cells assigned after re-performing clustering with only *E. coli* transcripts. **I.** Same as (G) but with color gradient indicating normalized λ phage UMI count in each cell. **J.** Normalized λ UMI count across each cluster in (H). **K.** Silhouette scores computed using the principal components of the lytic cluster (see panel D) and of “null subpopulation” which is a random sample of cells across each alignment. Comparison to (E) shows that the silhouette score of the lytic cluster drastically decreases with the removal of the λ genes

Using our M3-Seq data, we next sought to determine if *E. coli* mount an active transcriptional response to λ infection and lysis. Examining host genes that were differentially expressed between the lytic cluster and the rest of the population revealed only a small set of genes with modest log_2_ fold changes (<0.3 for upregulated genes) (Fig. 5F) and those that were upregulated were composed primarily of genes previously reported to be part of indirect effects (e.g., dNTP depletion from phage DNA replication or envelope stress from phage particles^32^). Reanalyzing our data using only the *E. coli* MG1655 genome next revealed that without inclusion of the phage genes, cells with high viral load could not be discriminated, i.e., we found that cell clusters determined by host transcriptional differences had similar average levels of λ transcripts (5% of the total UMIs, Fig. 5G-J). As another means of quantifying this observation, we used silhouette scores. Silhouette scores are a “goodness of clustering” metric that quantifies similarity of data within a cluster of interest compared to data outside that cluster^37^. We computed these scores for cells undergoing lysis (i.e., cells in cluster 3 from the combined analyses) using genes from both the *E. coli* and λ genome, genes from only the λ phage genome, and genes from only the *E. coli* genome (Fig. 5K). This analysis showed that cells undergoing lysis had a strong silhouette score when reads were aligned to λ genes, but not when aligned to only the *E. coli* genome. These results show that *E. coli* do not mount a specific, meaningful transcriptional response to λ phage lysis, despite the production of hundreds of foreign virions within the cell. We anticipate that this approach will be of great utility to the field of host-phage competition, especially in bacterial species which are not lab adapted and have active innate and adaptive immune systems.

## Discussion

While emerging technologies for scRNA-seq provide a means to identify and characterize rare subpopulations of bacteria, many meaningful applications will require the ability to sequence large numbers of single cells across a diverse array of experimental manipulations. Here, we report the development of M3-Seq, a two-step procedure of combinatorial indexing and *post hoc* ribosomal RNA depletion that simultaneously enables scale in the number of cells profiled (herein 229,671), breadth in the conditions that can be profiled in a single experiment (herein 20), and a high mRNA detection efficiency (herein 100-1,000 UMIs per cell) (Fig. 1A). Compared to existing methods for scRNA-seq in bacteria, M3-Seq holds advantages. Principally, the approach allows transcriptome-scale scRNA-seq at massive cell numbers while also removing abundant rRNA sequences—notably, while limiting enzymatic reactions on pre-amplified transcripts, each of which may add to loss of information. The approach can also be easily applied across multiple conditions. By contrast, pioneering combinatorial indexing-based methods provide reasonable scale in terms of the number of cells that can be profiled and comparable UMI capture but suffer from an overabundance of rRNA reads in the final library, which ultimately limits application^8,9^. Alternatively, probe-based approaches avoid signal from rRNA by design, but these approaches are not readily scaled to multiple conditions (i.e., probes are strain-specific)^11^ and, when the readout is reliant on imaging, may only be able to capture up to a hundred genes at a time^10^. Finally, we note that a recent preprint reports a method that combines many of the same elements of our approach, albeit without *post hoc* mRNA depletion and with more *in situ* enzymatic steps, possibly leading to a lower mRNA detection efficiency^38^. More broadly, though, we note that the many investments in technical development are indicative of the excitement in the field to push the envelope of scale and sensitivity.

Nevertheless, challenges remain. Unlike imaging-based methods, M3-Seq does not capture spatial information. The approach is thus not immediately applicable to studies of biofilms. Although, as a general point, scRNA-seq-based analysis of biofilms and mixed-species bacterial communities will also benefit from careful development of species-agnostic cell fixation and permeabilization procedures. Indeed, in one of our own experiments (BW4), we attempted to profile four species of bacteria (*B. subtilis, E. coli, Pseudomonas aeruginosa, Staphylococcus aureus*) but found that we could not recover UMIs at a satisfactory capture rate for each species, which we attribute to physical differences. The success of detecting multiple species in these experiments at all, though, provides solid precedent for what we anticipate will be many applications of M3-Seq to exploring new niches and single-cell strategies that emerge within a microbial community.

Why do rare bacterial subpopulations exist within a genetically identical bacterial population? One reason may be that transcriptional heterogeneity can act as a bet-hedging strategy in response to environmental variation. Such effects have been challenging to study with previous methods, but using M3-Seq, we discovered a rare acid-tolerant subpopulation expressing the *gad* genes in *E. coli*. Through genetic manipulation and orthogonal validation, we found that *gad*-expressing bacteria could survive strong acid treatment but were less fit in standard growth conditions. The rare *gad*-expressing population we observed at early stationary phase (i.e., before the environment may acidify) therefore supports a bet-hedging model of gene expression. Indicative of scRNA-seq as a discovery platform, many questions remain about this observation: what mechanisms generate this transcriptional heterogeneity? How do varying environments change the presence of this subpopulation? How prevalent is this bet-hedging strategy in nature?

Looking forward, we see multiple biological systems for which our technology is ripe to be applied. Undoubtedly, a key application will be host-pathogen interactions. When we infected *E. coli* with λ phage, we captured both pathogen (λ) and host (*E. coli*) mRNA transcripts in individual cells. We anticipate that applying our method in targeted experiments may provide new insights into how bacteria mobilize immunity mechanisms in response to phage infection. Moreover, this application need not be restricted to bacterial cells. Because of the generality of using random primers and the rRNA depletion scheme, our method can also be employed to study how mammalian cells respond to infection by intracellular pathogens, and how these infecting pathogens respond to host factors.

## Author contributions

B.W., B.S.A., N.S.W., and Z.G. were responsible for the conception, design, and interpretation of the experiments and wrote the manuscript with input from all authors. B.W. and A.E.L developed the *post hoc* rRNA depletion pipeline. B.W, A.E.L, J.Y., M.K. conducted experiments. B.W. performed data analysis.

## Acknowledgments

We thank Wei Wang and the Genomics Core Facility of the Lewis-Sigler Institute, the Adamson, Gitai, and Wingreen labs for input, and Yuri Pritykin and Ryan McNulty for critically reading and providing feedback on the manuscript. This work was supported by the National Science Foundation (Center for the Physics of Biological Function, PHY-1734030 to N.S.W.; NSF MCB-2033020 to Z.G.), the NIH (R01 GM082938 to N.S.W.; NIH DP1AI124669 to Z.G.; the Princeton QCB training grant, NIH T32HG003284), and the German Research Foundation (Award Ko5239/1-1 to M.D.K). J.Y. was supported by a fellowship provided by the China Scholarship Council (CSC), based on the April 2015 Memorandum of Understanding between the CSC and Princeton University. A.E.L. was supported by the Damon Runyon Cancer Research Foundation Postdoctoral Fellowship (DRG-2432-21).

## Data and materials availability

Sequencing data will be deposited on GEO; all analysis and demultiplexing scripts are uploaded https://github.com/brwaang55/m3seq_scripts. Any raw image files will be available upon request.

## Supplementary Materials

Figs. S1 to S8

Tables S1 to S6

Movies S1 to S2

## Materials and Methods

### Experimental methods

#### Bacterial strains and growth conditions for BW1

*B. subtilis* 168 and *E. coli* (MG1655) were streaked out from a frozen glycerol stock onto an LB plate and grown overnight at 37°C. Following a night of growth, a single colony was picked and inoculated into 5 mL of LB broth and grown shaking at 250 RPM overnight at 37°C. The next morning, the overnight culture was diluted (1:100 for *E. coli*, 1:25 for *B. subtilis*) into multiple tubes 5 mL of fresh LB media in a 30 mL tube grown shaking at 250 RPM. Cells were harvested once at OD=0.6, and again 4 hours post dilution. The volume of cells was normalized so that 1 OD of cells was sampled and fixed at each step. Cells were immediately spun down for 5 minutes at 5,000 g at 4°C, resuspended in 4 mL of freshly made 4% formaldehyde. The resuspended cells were rotated overnight at 4°C until the next morning.

#### Bacterial strains and growth conditions for BW2

*B. subtilis* 168 and *E. coli* (MG1655) were streaked out from a frozen glycerol stock onto an LB plate and grown overnight at 37°C. Following a night of growth, a single colony was picked and inoculated into 5 mL of LB broth and grown shaking at 250 RPM overnight at 37°C. The next morning, the overnight culture was diluted (1:100 for *E. coli*, 1:25 for *B. subtilis*) into 35 mL of fresh LB media in a 250mL Erlenmeyer flask and grown shaking at 250RPM. Upon reaching OD = 0.3, 5 mL of cells were split into tubes containing 2X the minimum inhibitory concentration of antibiotics (ciprofloxacin or cefazolin, 2 tubes), or no drug (2 tubes). The cells in the no drug tubes were sampled once at OD = 0.6, and again 120 minutes after the split. The cells in the tubes with drugs were sampled 20 minutes post-split (T20), and again at 120 minutes post-split (T360). The volume of cells was normalized so that 1 OD of cells was sampled and fixed at each step. Cells were immediately spun down for 5 minutes at 5,000g at 4°C, resuspended in 4 mL of freshly made 4% formaldehyde. The resuspended cells were rotated overnight at 4°C until the next morning.

#### Bacterial strains and growth conditions for BW3

*B. subtilis* 168 and *E. coli* (MG1655 and Nissle) were streaked out from a frozen glycerol stock onto an LB plate and grown overnight at 37°C. Following a night of growth, a single colony was picked and inoculated into 5 mL of LB broth and grown shaking at 250 RPM overnight at 37°C. The next morning, the overnight culture was diluted (1:100 for *E. coli*, 1:25 for *B. subtilis*) into 35 mL of fresh LB media in a 250mL Erlenmeyer flask and grown shaking at 250RPM. Upon reaching OD = 0.3, 5 mL of cells were split into tubes containing 2X the minimum inhibitory concentration of antibiotics (ciprofloxacin or cefazolin), or no drug. The cells in the no drug tubes were sampled once at OD = 0.6, and again 360 minutes after the split. The cells in the tubes with drugs were sampled 90 minutes post-split (T90), and again at 360 minutes post-split (T360). The volume of cells was normalized so that 1 OD of cells was sampled and fixed at each step. Cells were immediately spun down for 5 minutes at 5,000g at 4°C, resuspended in 4 mL of freshly made 4% formaldehyde. The resuspended cells were rotated overnight at 4°C until the next morning.

#### Bacterial strains and growth conditions for BW4

*B. subtilis* 168, *E. coli* MG1655, and *P. aeruginosa* PA14 were streaked out from a frozen glycerol stock onto an LB plate and grown overnight at 37°C. Following a night of growth, a single colony was picked and inoculated into 5 mL of LB broth and grown shaking at 250 RPM overnight at 37°C. The next morning, the overnight culture was diluted (1:100 for *E. coli*, 1:25 for *B. subtilis*, 1:50 for *P. aeruginosa*) into 35 mL of fresh LB media in a 250mL Erlenmeyer flask and grown shaking at 250 RPM. Upon reaching OD = 0.3, 4mL of cells were split into tubes containing 2X the minimum inhibitory concentration of antibiotics (gentamycin, tetracycline, erythromycin, chloramphenicol, cefazolin, cycloserine, ciprofloxacin, or nalidixic acid), *λ* phage at MOI=100 (for *E. coli*), or no drug. The cells in the tubes were sampled and had their absorbance read 90 minutes post-split (T90). The volume of cells was normalized so that 1 OD of cells was sampled and fixed at each step. Cells were then prepared in the same manner as with BW1,2,3.

#### Cell preparation

Following an overnight fixation, cells were prepared for scRNA-seq as previously described^8^. Briefly, cells were first spun down for 10 minutes at 5,000g at 4°C. Cells were then resuspended in 0.5 mL of PBS-RI, which comprises of PBS + 0.01 U/μL SUPERase-IN RNase Inhibitor (Invitrogen, AM2696). Cells were spun down again for 10 minutes at 5,000g at 4°C and resuspended in 300 μL of 1X PBS-RI, and 300 μL of 100% ethanol. Following the first permeabilization, cells were spun down for 8 minutes at 7,000g at 4°C, and washed twice with 200 μL of PBS-RI. After this final wash, cells were permeabilized by resuspension in 45 μL of 2.5mg/mL lysozyme solution dissolved in TEL-RI buffer, comprised of 100mM Tris pH 8.0, 50mM EDTA, 0.1U/μL SUPERase-IN RNase Inhibitor, and incubated at 30°C for 15 minutes. Cells were then spun down and washed twice in 100 μL of PBS-RI. After the final wash, cells were resuspended in 100 μL of 0.5X PBS-RI, and counted and examined with a hemocytometer (INCYTO DHC-S02).

#### Round-one indexing

Fixed and permeabilized cells were split into wells of a 96 well plate, each containing a single indexing primer (2.5 μL/well, 20μM). To each well, we added 312,500 cells, 0.25 μL of Maxima H Minus Reverse Transcriptase (Thermo Fisher Scientific, EP0753), 0.25 μL of dNTPs at an original concentration of 10 μM (NEB, N0447L), 2.5μL of 5X Maxima H Minus Reverse Transcription Buffer, 0.125 μL RNase-OUT (Thermo Fisher Scientific, 10777019), and PEG8000 to a final concentration of 7.5%, Tween-20 to a final concentration of 0.02%, and nuclease free water up to 10 μL. Reactions were then incubated as follows to perform first-round indexing by reverse transcription: 50°C for 10 minutes, 8°C for 12 seconds, 15°C for 45 seconds, 20°C for 45 seconds, 30°C for 40 seconds, 42°C for 6 minutes, 50°C for 50 minutes, and hold at 4°C. Samples were then pooled together and spun for 20 minutes at 7,000 g to isolate processed cells. Cells were then washed in 0.5 X PBS-RI and resuspended in 75 μL of 1X Ampligase buffer (Lucigen, A0102K). Pooled cells were counted and examined on the hemocytometer and diluted for loading onto the Chromium Controller (10x Genomics). The cell loading for each experiment indicated in Supplementary Table 2. Methods in this section were adapted from single-cell combinatorial fluidic indexing procedures.

#### Loading cells into microfluidic droplets

Cells were prepared for loading onto the Chromium scATAC platform v1.1 (10X Genomics 1000176). After counting, pooled cells were aliquoted with and mixed with 19 μL 10X Ampligase Buffer, 2.3μL U/μL Ampligase (Lucigen A0102K), 1.5 μL Reducing agent B (10x Genomics 2000087), 2.3 μL of 100 μM bridge oligo oDS025, and nuclease free water up to 75 μL. The mixture was kept on ice and loaded onto the Chromium Next GEM Chip H (10x Genomics, 1000162) with gel beads from the Chromium Next GEM Single Cell ATAC Library & Gel Bead Kit (10x Genomics, 1000176). To create emulsions, we followed the Chromium Single Cell ATAC Reagent Kits User Guide (v1.1 Chemistry) (CG000209 Rev A). Briefly, the microfluidic chip was prepared by adding 70μL of cell mixture to wells in row 1, 50μL Next GEM scATAC beads to wells in row 2, and 40μL of partitioning oil to wells in row 3. Additionally, 50% glycerol was added to all unused lanes (40μL of 50% glycerol was added to unused lanes in row 3, 50μL to unused lanes in row 2, and 70μL to unused lanes in row 1). The chip runs on the Chromium Controller (10x Genomics) with the Next GEM Chip H program. This step partitions the cells and uniquely indexed gel beads into droplets. Methods in this section were adapted from single-cell combinatorial fluidic indexing procedures^13^.

#### Round-two indexing

After transferring 100μL of each emulsion mixture to a clean reaction tube, second-round indexing was performed by ligation. Briefly, emulsions were incubated at 98°C for 30 seconds and 59°C for 2 minutes in 12 cycles. Emulsions were broken by adding 125μL Recovery Agent (10x Genomics) and pipetting up the hydrophobic phase. Cells were then reverse crosslinked and lysed by adding 10μL of 10X Lysis-T (250mM EDTA, 2M NaCl, 10% Triton X-100) and 4μL of proteinase K (NEB, P8107S) and incubating at 55°C for 1 hour. After lysis, DNA:RNA hybrid libraries were isolated with the following procedure: (1) 200μL of Dynabead Cleanup Mix, which consists of 182 μL Cleanup Buffer (10X Genomics, 2000088), 9μL Dynabeads MyOne Silane (Thermo Fisher Scientific, 37002D), 5μL Reducing Agent B (10X Genomics, no catalog no.) and 5μL of nuclease free water was added to each sample; (2) samples were mixed by pipetting (10x); (3) samples were incubated at room temperature for at least 10 minutes; (4) beads were isolated from samples using a magnetic stand and washed 2 times with 200μL 80% ethanol; (5) hybrid libraries were then eluted in 40μL of elution buffer (Qiagen, 19086).

#### Second strand cDNA synthesis

The eluted single stranded library was stripped of RNA by adding 2μL of RNase H (NEB M0297L), 4 μL of 10X RNase H buffer (NEB B0297S) and incubating for 30 minutes at 37°C. The reaction was purified with a 1.8X SPRI, where the final eluate volume was 25 μL. To perform second strand synthesis, we used a modified version from Hughes et al.^39^, where we added 8μL of 5X Maxima H-Reverse Transcription Buffer, 4 μL of 10 μM dNTP’s, 2.5 μL of Klenow Fragment (3’ -> 5’ exo -, NEB M0212L), 5 μL 50% PEG 8000, and 1.5 μL of 100 μM S^3 randomer (oBW140). The reaction was incubated at 37°C for 60 minutes, cleaned with a 1.8X SPRI, and eluted in 30 μL of Nuclease free water. The full length, double stranded library was amplified using PCR by adding 30 μL of 2X Q5 High Fidelity Master Mix (NEB M0492L), 0.4 μL of 100 μM oDS028, and 0.4 μL of 100 μM oBW170. We amplified the library using the following protocol: 98°C for 30 seconds, 14 cycles of 98°C for 20 seconds, 65°C for 30 seconds, 72°C for 3 minutes. Following the first round of PCR, the reaction was cleaned twice once using a 1.2X SPRI reaction, each time eluting in 40 μL. This was to ensure primer dimers were properly removed. The resulting samples were the gene expression (GEX) libraries.

#### Library fragmentation using Tn5 transposase

We prepared the following 5X Tn5 reaction buffer: 50mM TAPS (Sigma, T9659-100G), 25mM MgCl2. We assembled Nextera Read 2-only transposomes according to established protocols^13^. Briefly, 10 μL of 100 μM oDS029 and 10 μL of 100 μM oDS30 were mixed and annealed using the following temperature program: 95°C for 2 minutes, followed by a 0.1°C/second ramp down to 4°C. Annealed oligos were then diluted with 80 μL of nuclease free water (final concentration, 10 μM) and, after 10 μL of 100% glycerol was added to the mixture, 8μL of the oligo-glycerol sample was mixed with 2μL of EZ-Tn5 (Lucigen, TNP92110) and incubated at 25°C for 40 minutes. The resulting Read 2 transposomes were stored at −20°C.

After construction, gene expression libraries were quantified (Qubit HS dsDNA kit) and fragmented in multiple reactions with the following components: 10 ng gene expression library sample, 4 μL of 5x Tn5 buffer, 1μL of Read 2 transposome, and water up to 20 μL. Reactions were incubated at 55°C for 10 minutes and then inactivated with 1μL of 20% SDS at 55°C for 10 minutes. Following inactivation, reactions were purified using a 1.2X SPRI reaction (elution volume, 25 μL). The resulting samples were the fragmented GEX libraries.

#### Second library amplification and *in vitro* transcription

Fragmented GEX libraries were mixed with 25 μL of 2X Q5 Master Mix, 0.4 μL of 100 μM oBW170, 0.4 μL of 100 μM oBW168 and amplified using the following protocol: 72°C for 3 minutes, 98°C for 30 seconds, 9 cycles of 98°C for 10 seconds, 65°C for 30 seconds, 72°C for 30 seconds, a final incubation at 72°C for 5 minutes, and hold at 4°C. Resulting samples were purified with a 1.2X SPRI reaction (elution volume, 40 μL) and converted into RNA by *in vitro* transcription. Briefly, 100ng of amplified libraries were mixed with 8μL 5X Transcription Buffer (Thermo Fisher Scientific, EP0112), 6μL of 2.5 mM rNTPs (NEB, N0466L), 1.5 μL of T7 RNA Polymerase (Thermo Fisher Scientific, EP0112), and 1μL of RNase-Out. Reactions were incubated at 37°C for 2 hours, after which DNA templates were digested with 3μL DNase I (NEB, M0303L) and 3μL 10X DNase I buffer (NEB, B0303S) at 37°C for 15 minutes. RNA was purified using a 2X SPRI reaction (elution volume, 25 μL). These samples were the *in vitro* transcribed GEX libraries.

#### Ribosomal RNA depletion

To enrich for mRNA reads within a DNA library that was constructed using random priming, we developed an in-house approach to deplete ribosomal reads. Probes hybridizing to ribosomal RNA sequences of the bacterial species used in this study were designed (using the software designed by Huang et al.^17^) and multiple reactions containing were prepared as follows (using protocols adapted from^17^): 500 ng of in vitro transcribed RNA, 3 μg of rRNA probes, 0.6 μL 5 M NaCl, 1.5 μL 1M Tris-HCl, and Nuclease free water up to 15 μL. Hybridization was then performed using the following temperature program: 95°C for 2 minutes, and 0.1°C/second ramp down to 25°C, 25°C for 5 minutes. Following rRNA probe hybridization, 6μL RNase H mix, which consists of 3μL of 10x RNase H buffer (NEB B0297), 2μL of Thermostable RNase H (NEB M0523S), and 1μL of RNase H were added to each tube. The reactions were incubated for 45 minutes at 50°C to digest the rRNA-DNA hybrids. Following rRNA digestion, the DNA probes were degraded by adding 3μL of 10x DNase I buffer, 3μL of DNase I, and incubating for 45 minutes at 37°C. The rRNA-depleted RNA library was purified with a 2x SPRI reaction and eluted in 25 μL of nuclease free water.

#### Final library prep

To recover an rRNA-depleted cDNA library for sequencing, we next performed a second round of reverse transcription using the end specific P5 primer, thus ensuring reverse transcription of full library constructs. To each tube of purified RNA, we added the following reagents: 8 μL Maxima H Minus Reverse Transcription Buffer, 1μL Maxima H Minus Reverse Transcriptase, 1μL RNase Out, 6μL 2.5mM dNTPs, 0.4 μL 100 μM oBW170, 0.2μL 100 μM oBW171. The reaction was incubated in the thermocycler with the following temperature program: 50°C for 10 minutes, 8°C for 12 seconds, 15°C for 45 seconds, 20°C for 45 seconds, 30°C for 40 seconds, 42°C for 6 minutes, 50°C for 18 minutes, and hold at 4°C.

Following reverse transcription, the reaction was purified with a 1.2X SPRI and eluted in 25 μL of nuclease free water. The reverse transcribed DNA reactions were then indexed using a final indexing PCR to multiplex different libraries on the same sequencing run. For each reaction, 25 μL of reverse transcribed DNA was mixed with 25 μL Q5 High Fidelity Master Mix, 0.4 μL 100 μM oBW501, and 0.4 μL 100 μM of a unique P7 index primer. The reactions were amplified with the following temperature program: 98°C for 30 seconds, 9 cycles of 98°C for 10 seconds, 65°C for 30 seconds, 72°C for 30 seconds, a final incubation at 72°C for 5 minutes, and hold at 4°C.

After two purifications with 0.8X SPRI, our final sequencing libraries were quality controlled on the Qubit and Bioanalyzer. We also checked the concentration and quality of each DNA library using qPCR (primers: oBW170/oBW176, oBW141/oBW176). We note that this final qPCR step is essential as it checks for the percentage of the reads that can be sequenced in each library. Typically, a ΔCT of 0-0.6 (oBW141/oBW176 - oBW170/oBW176) indicates a fully sequenceable library. Following the final qPCR, libraries were diluted to 5nM, and sequenced with the NovaSeq SP 100 cycle kit (Illumina 20028401) using the following read structure: 26bp Read 1, 30bp i5 index, 8bp i7 index, 74bp Read2.

#### Fluorescent in-situ hybridization (FISH)

To enable cost effective detection of multiple different RNAs in the same cells, we closely followed established frameworks for single molecule FISH^40,41^. Briefly, multiple primary probes hybridizing to an mRNA of interest are first designed. These probes contain a constant 20nt flanking sequence that allows for hybridization of a fluorescent secondary probe. This allows us to avoid the cost of ordering multiple fluorescent primary probes to tile our gene of interest.

Primary probes for fluorescent in-situ hybridization for RNA sequences of interest were designed using the same software used to design rRNA probes^17^. For each RNA transcript of interest, we designed at least 10 different probes hybridizing to different regions of that transcript. A 20nt sequence was added to the 3’ end of each probe to allow for hybridization of the fluorescent readout probes. Primary probes for each gene were mixed at an equimolar ratio such that the final concentration of DNA molecules was 100 μM. Fluorescent readout probes were ordered as previously described^41^.

Cells in each condition of interest were grown, fixed, and permeabilized as described above. After the permeabilization step, cells were washed and resuspended in 600 μL primary hybridization buffer (40% Formamide (Thermo Fisher Scientific 15515026), 2X SSC (Invitrogen AM9673)) and aliquoted into 1.5 mL tubes. 1μL of 100 μM primary probe mix was added to each tube and hybridized overnight at 30°C in the dark. The next morning, cells were spun down at 7,000g for 8 minutes and resuspended in 200 μL wash buffer (30% Formamide (Thermo Fisher Scientific 15515026), 2X SSC (Invitrogen AM9673)). Cells were spun down for 8 minutes at 7,000g, resuspended again in 200 μL wash buffer, and incubated in the dark at room temperature for 30 minutes. Cells were then spun down at 7,000g for 8 minutes and resuspended in 100 μL secondary hybridization buffer (10% Formamide, 2X SSC, 10% Ficoll PM-400 (Sigma-Aldrich F5415-25 mL)). 0.5 μL of each 100 μM readout probe was added to the tubes, and incubated for 1 hour at 34°C. Following secondary hybridization, cells were spun down at 7,000g, and resuspended in wash buffer with 10 μg/mL DAPI (Thermo Fisher Scientific D1306). Cells were incubated for 20 minutes at room temperature, spun down at 7,000g, and resuspended in 100 μL of 2X SSC.

Cells were imaged on 1% agarose pads made with filtered water on a Nikon TiE microscope with a Plan Apo 100X objective, and Hanamatsu ORCAFlash4.0 camera. Images were analyzed using FIJI software.

#### Acid tolerance assay

A 25mL culture of *E. coli* (MG1655) or *E. coli* (MG1655 *ΔgadAΔgadBΔgadC*) was first grown to OD = 0.3 in a 125mL flask shaking at 250 RPM 37°C. After reaching OD = 0.3, the cultures were split in aliquots of 5mL to culture tubes and placed back onto the shaker to grow for another 6 hours until OD = 2.8. Cultures were then acidified to pH 3.0 using 12N HCl and returned to the shaker. 10 μL of the cultures was sampled at intermittent timepoints and serial diluted for CFU counting.

#### Acid recovery assay

A 25mL culture of *E. coli* (MG1655) transformed with P_*gadB*_-GFP was first grown to OD = 0.3 in a 125mL flask shaking at 250 RPM 37°C. After reaching OD = 0.3, the cultures were split in aliquots of 5mL to culture tubes and placed back onto the shaker to grow for another 6 hours until OD = 2.8. At this point, 1 μL of the culture was imaged on an 1% agarose pad made with LB media to understand the distribution of GFP fluorescence in single cells. Cultures were then acidified to pH 3.0 using 12N HCl and returned to the shaker. Following an hour of acid stress, 1 μL of the acidified culture was transferred onto an 1% agarose pad made with fresh LB media to assess viability. Cells were imaged every 15 minutes to track and assess growth over time.

The resulting movies were analyzed by first segmenting the cells using Delta^42^, and then using custom python scripts to extract the fluorescence distribution and assess viability. A cell was considered viable if it underwent a single division during the 8-hour imaging period.

#### Bulk RNA-seq Library preparation

*E. coli* (MG1655) was grown as described above to OD = 0.6. 2mL of cells were spun down at 5,000g for 10 minutes, resuspended in 45 μL of 2.5mg/mL lysozyme solution (described above), and incubated at 37°C for 15 minutes. RNA was purified using the Qiagen RNeasy Mini Kit (Qiagen 74104) where the final eluate volume was 30 μL. The RNA was reverse transcribed by adding 5 μL Maxima H Minus Reverse Transcription Buffer, 0.5 μL Maxima H Minus Reverse Transcriptase, 0.5 μL RNase Out, 4 μL 2.5mM dNTPs, 0.4 μL 100 μM oBW 121 and incubating using the following temperature program: 50°C for 10 minutes, 8°C for 12 seconds, 15°C for 45 seconds, 20°C for 45 seconds, 30°C for 40 seconds, 42°C for 6 minutes, 50°C for 50 minutes, and hold at 4°C.

Following reverse transcription, RNA was stripped from the reverse transcribed DNA by adding 2μL of RNase H and incubating the mixture at 37°C for another 30 minutes. The library was purified using a 1.2X SPRI and eluted in 25 μL nuclease free water. Second strand synthesis, PCR, and tagmentation were performed as described above. The first PCR was performed using primer pairs oBW154 and oDS28. Following tagmentation, the library was amplified 8 cycles as described above using oBW154 and oBW168. This library was used to test for different rRNA depletion strategies.

#### Cas9 based rRNA depletion

To test Cas9 based rRNA depletion, we first synthesized a pool of guide RNAs which cleave at different sites of the 5S, 16S, and 23S ribosomal RNAs. DNA templates for the guide RNAs were designed using previously described software^15^. The DNA templates were purchased as a pool from IDT, and amplified with PCR by first annealing at a 1:1 equimolar ratio mixing 1μL DNA template, 0.4 μL 100 μM oBW138, 0.4 μL 100 μM oBW139, 10 μL nuclease free water, 12.5 μL 2X Q5 High Fidelity Master Mix, and using the following temperature program: 98°C for 30 seconds, 35 cycles of 98°C for 10 seconds, 65°C for 30 seconds, 72°C for 45 seconds, a final incubation at 72°C for 5 minutes, and hold at 4°C. Following PCR, the DNA templates were purified using a 1.2X SPRI and used for in vitro transcription. Guide RNAs were transcribed using the NEB HiScribe kit (NEB E2040S) by mixing 100ng of DNA template, 2μL of 10X reaction buffer, 2μL 100mM ATP, 2μL 100mM GTP, 2μL 100mM CTP, 2μL 100mM UTP, 2μL T7 RNA Polymerase Mix, nuclease free water up to 20 μL, and incubated overnight at 37°C.

Following an overnight in vitro transcription, DNA template was digested by adding 3μL 10X DNase buffer, 2μL DNase I, and incubating for an additional 15 minutes at 37°C. Guide RNAs were purified using a 2X SPRI reaction and checked for purity by running on a 15% TBE-Urea Gel (Invitrogen EC6885BOX). Guide RNA concentration was quantified using the Broad Range RNA Qubit kit (Thermo Fisher Scientific Q10210).

To perform Cas9 based depletion in our most optimized condition, 2 ng of library was mixed with 1.5 μL NEB 3.1 buffer, and sgRNA and NEB cas9 at a 20,000:3,000:1 ratio of sgRNA:Cas9: DNA. The reaction was incubated at 37°C for 2 hours after which Cas9 was stripped from the DNA by adding in 1μL Proteinase K, 1μL 10% SDS, and incubating for 15 minutes at 50°C. The DNA library was purified with a 1.2X SPRI, eluted in 25 μL nuclease free water, and mixed with 25 μL 2X Q5 High Fidelity Master Mix, 0.4 μL 100 μM oBW170, and 0.4 μL 100 μM of a unique P7 index primer. The reactions were amplified with the following temperature program: 98°C for 30 seconds, 12 cycles of 98°C for 10 seconds, 65°C for 30 seconds, 72°C for 30 seconds, a final incubation at 72°C for 5 minutes, and hold at 4°C. Libraries were sequenced on the MiSeq Reagent Kit v2 (300 cycles) (Illumina MS-102-2002) using the following read structure: 26bp Read 1, 30bp i5 index, 8bp i7 index, 100bp Read2.

#### Quantifying cell loading in the 10X Microfluidic system

To quantify if single bacterial cells could be loaded into the 10X Microfluidic system, we first fixed 2mL of *E. coli* MG1655 cells grown to OD=0.4 overnight in 4 mL of 4% formaldehyde. Cells were prepared as described above up to after the first wash following permeabilization. Following the first wash, cells were incubated in 50 μL of 5 μM Sytox Green (Thermo Fisher Scientific S7020) for 15 minutes. After the incubation, cells were washed twice in 100 μL of PBS-RI, and then resuspended in 100 μL of 0.5X PBS-RI. Cells were counted, and then loaded onto the 10X Microfluidic system using the Chip A 5’ kit.

Following droplet generation, 5uL of the mixture was transferred onto a glass coverslip and imaged on a Nikon TiE microscope with a Plan Apo 20X objective, and Hanamatsu ORCAFlash4.0 camera. Cells in each droplet were then manually counted.

### Computational Methods

#### Data preprocessing

Raw basecalls were retrieved from the NovaSeq, and processed with a custom version of picard tools v2.19.2 following the pipeline described in the original SciFi-Seq pipeline^13^. Reads were aligned to a combination of one or more of *B. subtilis* 168, *E. coli* MG1655, and *E. coli* Nissle genomes using STAR v2.76^43^ and annotated with featureCounts v2.0.0^44^. Reads were filtered such that all the reads used for downstream analysis have mapQ score > 1, which correspond to reads that have aligned to 3 or less locations, and mapped lengths greater than 20bp. Annotated and filtered reads were loaded into Python 3.7.6 where custom code was written to assign non-rRNA reads to combinations of droplet and plate barcodes in pandas.

After assigning reads to barcode combinations, we filtered out “cell clumps”, which we defined as droplet barcodes in which a given droplet barcode had more than 8 associated plate barcodes. We split barcode combinations by condition (round-one barcodes) and performed another filtering step using the knee method for each condition^4,8^. We note that this step is important because bacteria in different conditions have different amounts of mean mRNA expression. When necessary, index collision rates were calculated by computing the fraction of cells with <85% of UMIs assigned to one species, and then correcting the collision rate using previously described methods^14^. After the last filtering step, a cell/gene matrix was made where the entries of the matrix are the number of UMIs that we measured for that gene in a particular cell.

#### Single-cell analysis

Metrics for the scRNA-seq results were compiled and plotted using custom scripts in Python 3.7.6. Downstream analysis of single-cell data was performed using pipelines detailed in Seurat v4.0.3^45^. Data were first preprocessed by filtering out genes that were expressed in less than 10 cells and cells that expressed less than 10 UMIs. The data were then normalized by dividing the UMI counts in each cell by the total number of UMIs measured in that cell, multiplying by a scale factor of 100, adding a count of 1 to each entry, and then lognormalizing the scaled values^45^. The normalized expression data were then scaled to have mean 0 and unit variance, and dimensionally reduced using principal component analysis. When necessary, the kurtosis of each principal component was computed by taking the matrix of cells by principal component coordinates and then calling the “kurtosis” function from the R package moments^46^.

Following principal component analysis, we computed a uniform manifold approximation representation and a shared neighbor graph (SNN) using the first 10 principal components. We performed graph-based clustering on the shared neighbor graph to identify clusters of gene expression programs using the Louvain algorithm (algorithm 3 in Seurat 4.0.3). Marker genes for each cluster were computed using the Wilcoxon Rank-sum test. Further data analysis and plotting was performed using custom scripts in R.

Gene set enrichment analyses were performed using topGo (2.48.0). Briefly, marker genes were determined using the FindMarkers function in Seurat, where we by compared the within-cluster average expression to out of cluster average expression and filtering for genes with *p*-value < 0.05 (Wilcoxon-rank sum test). This list was then split into genes that were upregulated in the cluster and genes that were downregulated. The two lists of genes were then used for biological process term enrichment using Fisher’s exact test, in which the input was a vector of length (number of genes in the genome), and each entry in the vector was 1 if the index corresponded to a gene in the list of upregulated/downregulated (depending on if we were testing up- or downregulated genes) genes and 0 otherwise. Following the test, the *p*-values are -log_10_ transformed such that the most strongly enriched biological processes have the highest score. Selected processes to be plotted were those with the lowest *p*-values after thresholding at 0.05.

To compute silhouette scores, we took the PCA matrix and cluster outputs from Seurat, and used the silhouette score function from the KBET package^47^.

#### Comparison with bulk RNA-seq

Bulk RNA-seq data^8^ for exponentially growing *E. coli* were obtained from Blattman et al. (GEO accession number GSE141018). Raw reads from the bulk data were aligned to the *E. coli* MG1655 genome and annotated as described above. Single-cell and bulk transcriptomes of exponential growing *E. coli* were compared by computing the Pearson correlation of log_10_ transcripts per million (TPM) of each gene between the two measurements. TPM for each gene in single-cell data was then computed by adding a pseudocount of 1 to each gene, summing over the UMI counts for that gene across all cells, normalizing by gene length, and dividing by the sum of length normalized counts. TPM for bulk measurements were computed as previously described. The TPMs of the bulk and single-cell datasets were log_10_ transformed and used for plotting and correlation measurements.

