## Supplementary files for "Massively-parallel Microbial mRNA Sequencing (M3-Seq) reveals heterogeneous behaviors in bacteria at single-cell resolution"

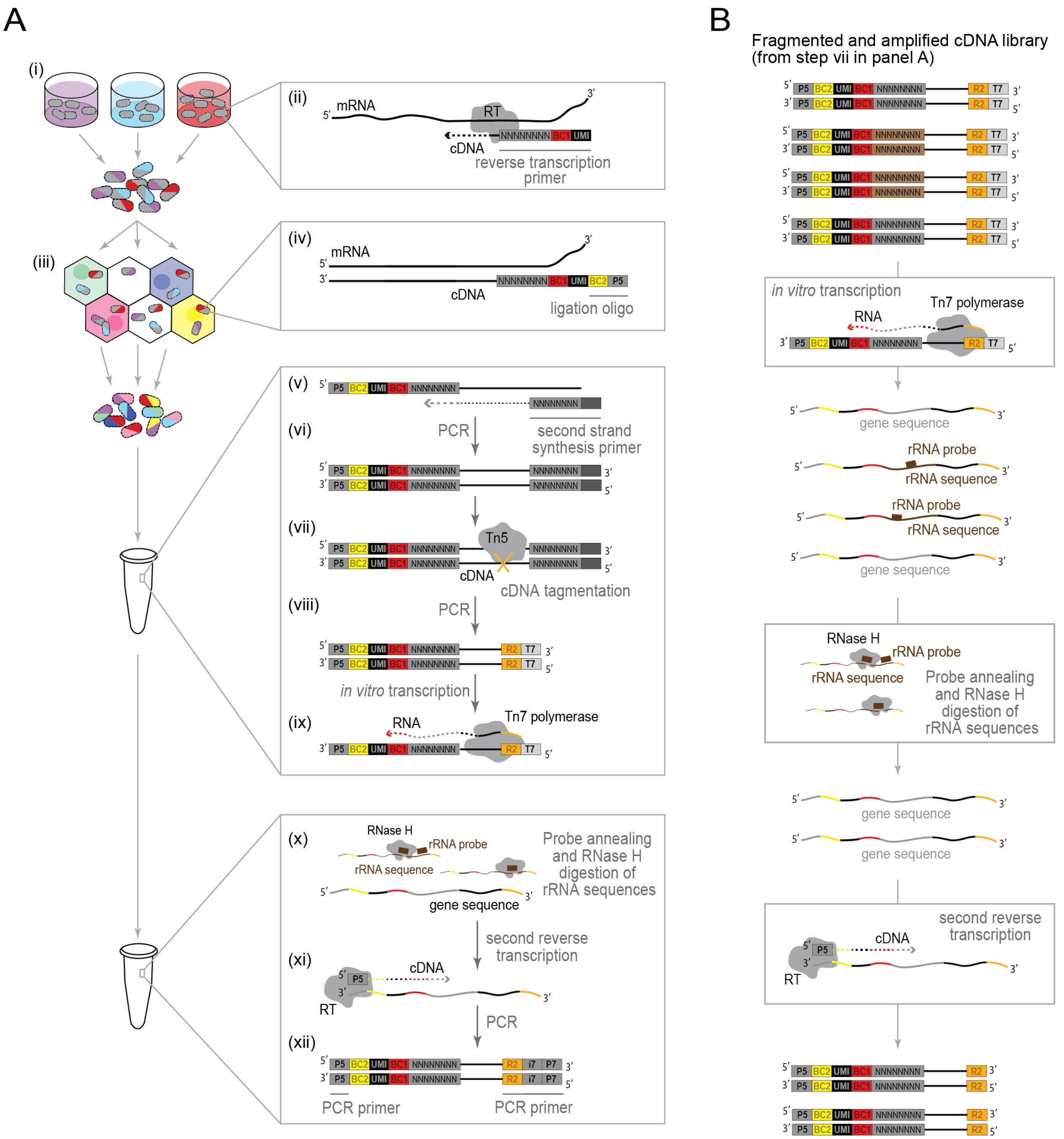
A
B
Fragmented and amplified cDNA library
(from step vii in panel A)
(i)
(ii)
mRNA
RT
cDNA
reverse transcription primer
(iii)
(iv)
ligation oligo
(v)
(vi)
PCR
second strand synthesis primer
(vii)
Tn5
cDNA tagmentation
(viii)
in vitro transcription
(ix)
RNA
Tn7 polymerase
(x)
RNase H
rRNA probe
rRNA sequence
gene sequence
Probe annealing and RNase H digestion of rRNA sequences
(xi)
second reverse transcription
(xii)
PCR primer
P5
BC2
UMI
BC1
NNNNNNNN
R2
T7
i7
P7

**Figure S1. M3-Seq experimental workflow and rRNA depletion scheme. A.** Detailed schematic of M3-Seq experimental workflow: Populations of fixed and permeabilized bacteria are (i) aliquoted into wells of one or more 96 well plates. Each well contains a uniquely indexed random hexamer, which acts as a primer for (ii) *in situ* reverse transcription. These primers also carry unique molecular identifier (UMIs) sequences. During reverse transcription, cell-associated RNA molecules are converted to cDNAs with primer barcodes and UMIs on their 5' ends. After reverse transcription, (iii) cells are pooled and loaded into a commercially available device for droplet-based indexing (herein, the Chromium Controller from 10x Genomics) without a need for limiting dilution. After partitioning into droplets, (iv) a second index is ligated onto the 5' end of the reverse transcribed, cell-associated cDNAs (herein, using Next GEM Single Cell ATAC reagents from 10x Genomics). Following indexing, cells are lysed, and (v) cDNA molecules are converted to double-strand DNA using a Klenow enzyme and a random primer with a PCR handle at the 5' end. This double-strand cDNA is then (vi) amplified by PCR, (vii) fragmented with Tn5 transposase loaded with Nextera read 2 primers, and (viii) attached to a T7 promoter via a second round of PCR. Next, (ix) cDNA molecules are transcribed back into RNA using T7 RNA polymerase. This step prepares the amplified library for rRNA depletion. After transcription, (x) the resulting RNA is annealed to a set of DNA probes that are complementary to rRNA sequences within the library (Table S4). This annealing allows for selective degradation of those sequences with RNase H. Finally, in a second reverse transcription step, (xi) the indexed and rRNA-depleted library is converted back into cDNA, and (xii) the resulting cDNA is amplified one more time to add a required sequencing adaptor. The library is then ready for paired end sequencing. **B.** Detailed schematic of rRNA depletion steps: To remove rRNA sequences from M3-Seq libraries, we (i) convert indexed and amplified cDNA libraries into RNA via *in vitro* transcription, (ii) hybridized rRNA sequences within the library to DNA probes and digest those sequences using RNase H, and (iii) convert the remaining sequences back into DNA using a P5 primer.

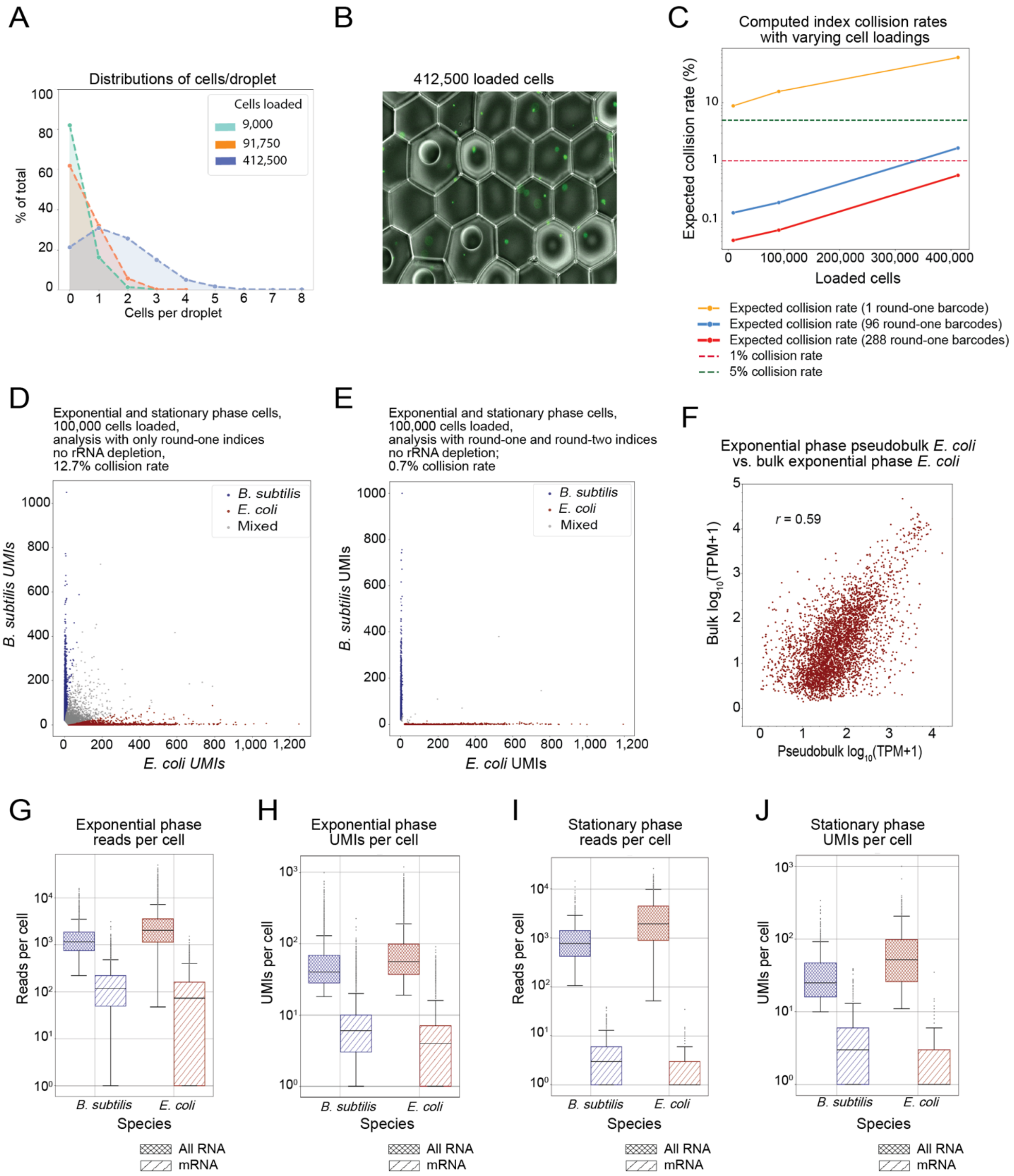
A
Distributions of cells/droplet
Cells loaded
9,000
91,750
412,500
% of total
Cells per droplet
B
412,500 loaded cells
C
Computed index collision rates with varying cell loadings
Expected collision rate (%)
Loaded cells
Expected collision rate (1 round-one barcode)
Expected collision rate (96 round-one barcodes)
Expected collision rate (288 round-one barcodes)
1% collision rate
5% collision rate
D
Exponential and stationary phase cells, 100,000 cells loaded, analysis with only round-one indices no rRNA depletion, 12.7% collision rate
B. subtilis
E. coli
Mixed
B. subtilis UMIs
E. coli UMIs
E
Exponential and stationary phase cells, 100,000 cells loaded, analysis with round-one and round-two indices no rRNA depletion; 0.7% collision rate
F
Exponential phase pseudobulk E. coli vs. bulk exponential phase E. coli
r = 0.59
Bulk log10(TPM+1)
Pseudobulk log10(TPM+1)
G
Exponential phase reads per cell
Reads per cell
H
Exponential phase UMIs per cell
UMIs per cell
I
Stationary phase reads per cell
J
Stationary phase UMIs per cell
Species
All RNA
mRNA

**Figure S2. Piloting single-cell RNA-sequencing in bacteria without rRNA depletion. A.** Distributions represent bacterial cells per droplet produced on the Chromium Controller (10x Genomics) at indicated cell numbers. Data calculated from observed rates by counting the number of fixed and stained *E. coli* cells in each droplet across the three cell loading conditions. **B.** Representative image of droplets quantified in panel (A). *E. coli* cells, which were stained with Sytox green, are visible as green dots. **C.** Data represent expected index collision rates (i.e., percentage of cells with the same round-one index labeled with the same round-two index) as a function of loaded cells using different indexing schemes. The lowest number of cells plotted for each condition is within the recommended loading range (9,000 cells) and predicts a >10% collision rate without a round-one (plate-based) indexing step. In contrast, ~300,000 cells should yield <1% collision (dashed red) with only 288 round-one indices (solid red). **D.** Analysis of a mixture of exponential and stationary phase *B. subtilis* (blue) and *E. coli* (red) using only round-two (droplet-based) indexes. Species assignments for each "cell" were made if >85% of corresponding UMIs mapped to species-specific transcripts. Otherwise, cells were designated as mixed. Data were generated without rRNA depletion (BW1 in Table S2). **E.** Same as (D) but analysis performed with combinatorial barcodes (i.e., both round-one and round-two indexes). Species assignments for each "cell" were made if >85% of corresponding UMIs mapped to species-specific transcripts. Otherwise, cells were designated as mixed. **F.** Comparison of published RNA-seq data[8] to computationally rRNA-depleted single-cell gene expression in exponential phase *E. coli* from BW1. Pseudobulk measurements were obtained by converting UMI counts to transcripts per million (TPM). Each point represents a single gene. *r*, Pearson correlation. **G.** Read counts per cell from single-cell gene expression data without rRNA depletion (median of 1146 ± 475 reads for *B. subtilis* and 2021 ± 1072 reads for *E. coli* when considering all RNAs; median of 117 ± 81 reads for *B. subtilis* and 72 ± 72 reads for *E. coli* when considering only mRNAs). **H.** Same as (G) but for UMI counts per cell (median of 39 ± 15 UMIs for *B. subtilis* and 55 ± 24 UMIs for *E. coli* when considering all RNAs; median of 5 ± 5 UMIs for *B. subtilis* and 3 ± 3 UMIs for *E. coli* when considering only mRNAs). I. Same as (G) but for stationary phase *B. subtilis* and *E. coli* (median of 772 ± 414 reads for *B. subtilis* and 1944 ± 1307 reads for *E. coli* when considering all RNAs; median of 23 ± 23 reads for *B. subtilis* and 0 ± 0 reads for *E. coli* when considering only mRNAs). J. Same as (H) but for stationary phase *B. subtilis* and *E. coli* (median of 24 ± 11 UMIs for *B. subtilis* and 51 ± 31 UMIs for *E. coli* when considering all RNAs; median of 2 ± 2 UMIs for *B. subtilis,* and 0 ± 0 UMIs for *E. coli* when considering only mRNAs).

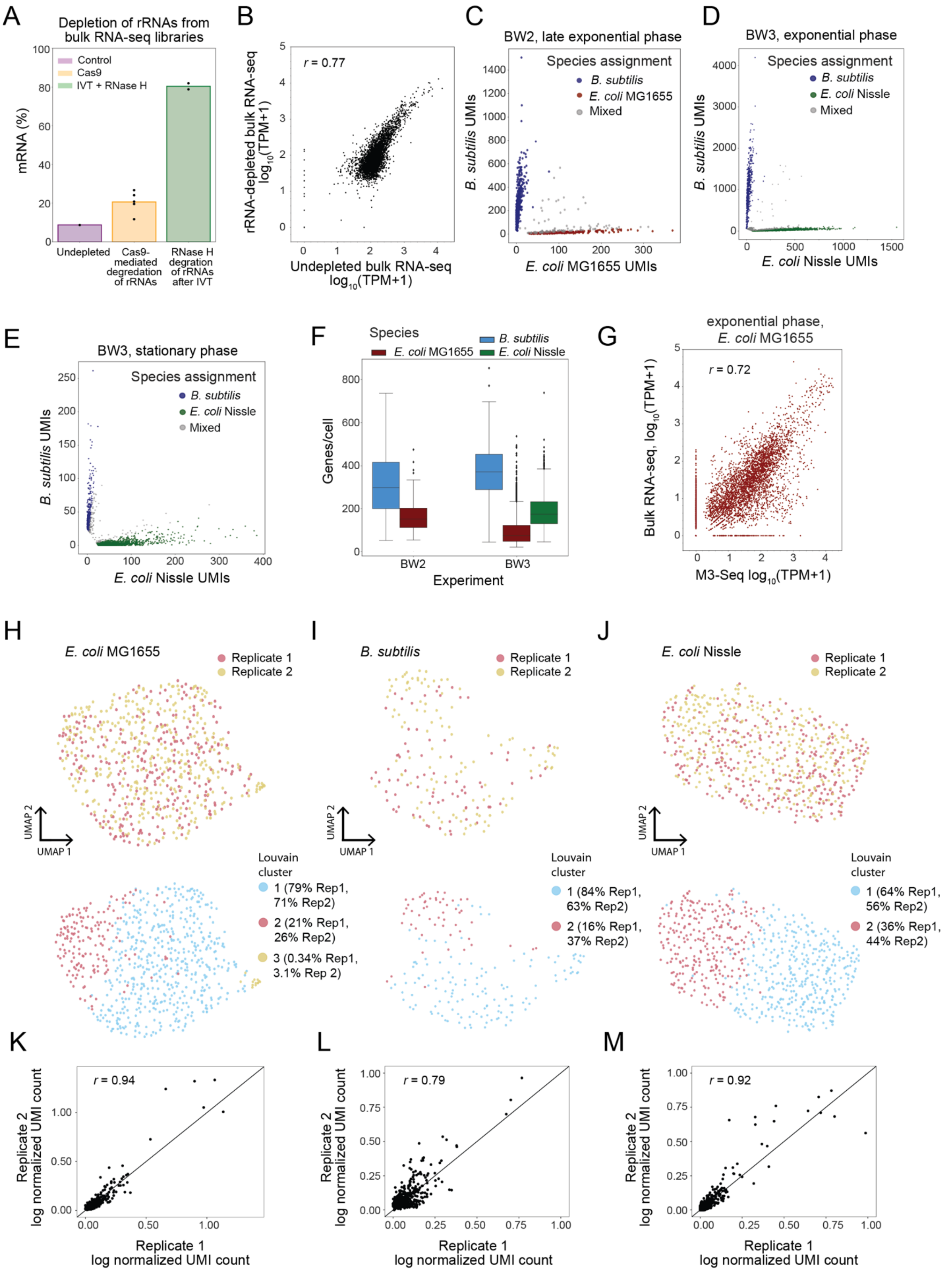
A
Depletion of rRNAs from bulk RNA-seq libraries
Control
Cas9
IVT + RNase H
mRNA (%)
Undepleted
Cas9-mediated degredation of rRNAs
RNase H degration of rRNAs after IVT
B
r = 0.77
rRNA-depleted bulk RNA-seq log10(TPM+1)
Undepleted bulk RNA-seq log10(TPM+1)
C
BW2, late exponential phase
Species assignment
B. subtilis
E. coli MG1655
Mixed
B. subtilis UMIs
E. coli MG1655 UMIs
D
BW3, exponential phase
Species assignment
B. subtilis
E. coli Nissle
Mixed
B. subtilis UMIs
E. coli Nissle UMIs
E
BW3, stationary phase
Species assignment
B. subtilis
E. coli Nissle
Mixed
B. subtilis UMIs
E. coli Nissle UMIs
F
Species
B. subtilis
E. coli MG1655
E. coli Nissle
Genes/cell
BW2
BW3
Experiment
G
exponential phase, E. coli MG1655
r = 0.72
Bulk RNA-seq, log10(TPM+1)
M3-Seq log10(TPM+1)
H
E. coli MG1655
Replicate 1
Replicate 2
UMAP 2
UMAP 1
Louvain cluster
1 (79% Rep1, 71% Rep2)
2 (21% Rep1, 26% Rep2)
3 (0.34% Rep1, 3.1% Rep 2)
I
B. subtilis
Replicate 1
Replicate 2
UMAP 2
UMAP 1
Louvain cluster
1 (84% Rep1, 63% Rep2)
2 (16% Rep1, 37% Rep2)
J
E. coli Nissle
Replicate 1
Replicate 2
UMAP 2
UMAP 1
Louvain cluster
1 (64% Rep1, 56% Rep2)
2 (36% Rep1, 44% Rep2)
K
r = 0.94
Replicate 2 log normalized UMI count
Replicate 1 log normalized UMI count
L
r = 0.79
Replicate 2 log normalized UMI count
Replicate 1 log normalized UMI count
M
r = 0.92
Replicate 2 log normalized UMI count
Replicate 1 log normalized UMI count

**Figure S3. Additional analysis of M3-Seq development. A.** Efficiency of rRNA depletion using two different *post hoc* approaches: degradation by rRNA-targeted Cas9[14,15] (yellow) and RNase H-mediated digestion after *in vitro* transcription (green). Without digestion roughly 10% of all reads within a bulk RNA-seq library align to mRNA. The former method improved this metric to roughly 20%, while the latter yielded 80% mRNA reads. **B.** Comparison of gene expression data from rRNA-depleted and control libraries produced. Bulk libraries were prepared as in the *Materials and Methods*. *r*, Pearson correlation. **C.** M3-Seq analysis of a mixture of *B. subtilis* (blue) and *E. coli* (red) in late exponential phase (OD = 2.1, 2.0 respectively) wherein each point corresponds to a single "cell" (i.e., unique combination of plate and droplet barcodes). Species assignments were made if >85% of corresponding UMIs mapped to unique species-specific transcripts. Otherwise, cells were designated as mixed (13% collision rate, 26% corrected). Data were generated with rRNA depletion (BW2 in Table S2). **D.** Same as (C) but for *B. subtilis* and a different strain of *E. coli* (OD = 0.3, 0.3 respectively). Data were generated with rRNA depletion (BW3 in Table S2) and show a 12% collision rate, 27% corrected. **E.** Same as (D) but for cells in stationary phase (OD = 2.4, 3.0 respectively). Data were generated with rRNA depletion (BW3 in Table S2) and show a 6.1% collision rate, 19% corrected. **F.** Genes per cell (after species assignment) observed in exponential phase cells across two experiments, BW2 and BW3 (298 ± 104 and 371± 82 median genes with absolute deviation for *B. subtilis*, respectively; 151 ± 47 and 75±31 median genes with absolute deviation for *E. coli* MG1655, respectively; 175 ± 50 genes with for *E. coli* Nissle in BW3). **G.** Comparison of published RNA-seq data[8] to M3-seq pseudobulk profiles from exponential phase *E. coli* from BW3. Pseudobulk measurements were obtained by converting UMI counts to transcripts per million (TPM). Each point represents a single gene. Note, in each experiment, *E. coli* were grown in different media (i.e., LB broth or EZ-Rich media). *r*, Pearson correlation. **H.** Top: UMAP projection of replicate M3-Seq data generated from *E. coli* MG1655 treated with twice the minimum inhibitory concentration of ciprofloxacin, sampled after 6 hours of treatment. Bottom: UMAP projection of replicate M3-Seq data with Louvain clusters overlaid. Within the cluster label is the percentage of cells (with respect to the total number of cells within replicate) belonging to that cluster from each replicate. **I.** Same as (H) but for *B. subtilis* 168. **J.** Same as (H) but for *E. coli* Nissle. **K.** Comparison of replicate data using mean log normalized UMI counts per cell (i.e., unique UMIs relative to total UMIs per cell averaged across all cells for each gene). Each point represents a single gene. *r*, Pearson correlation. **L.** Same as (K) but for *B. subtilis* 168. **M.** Same as (K) but for *E. coli* Nissle.

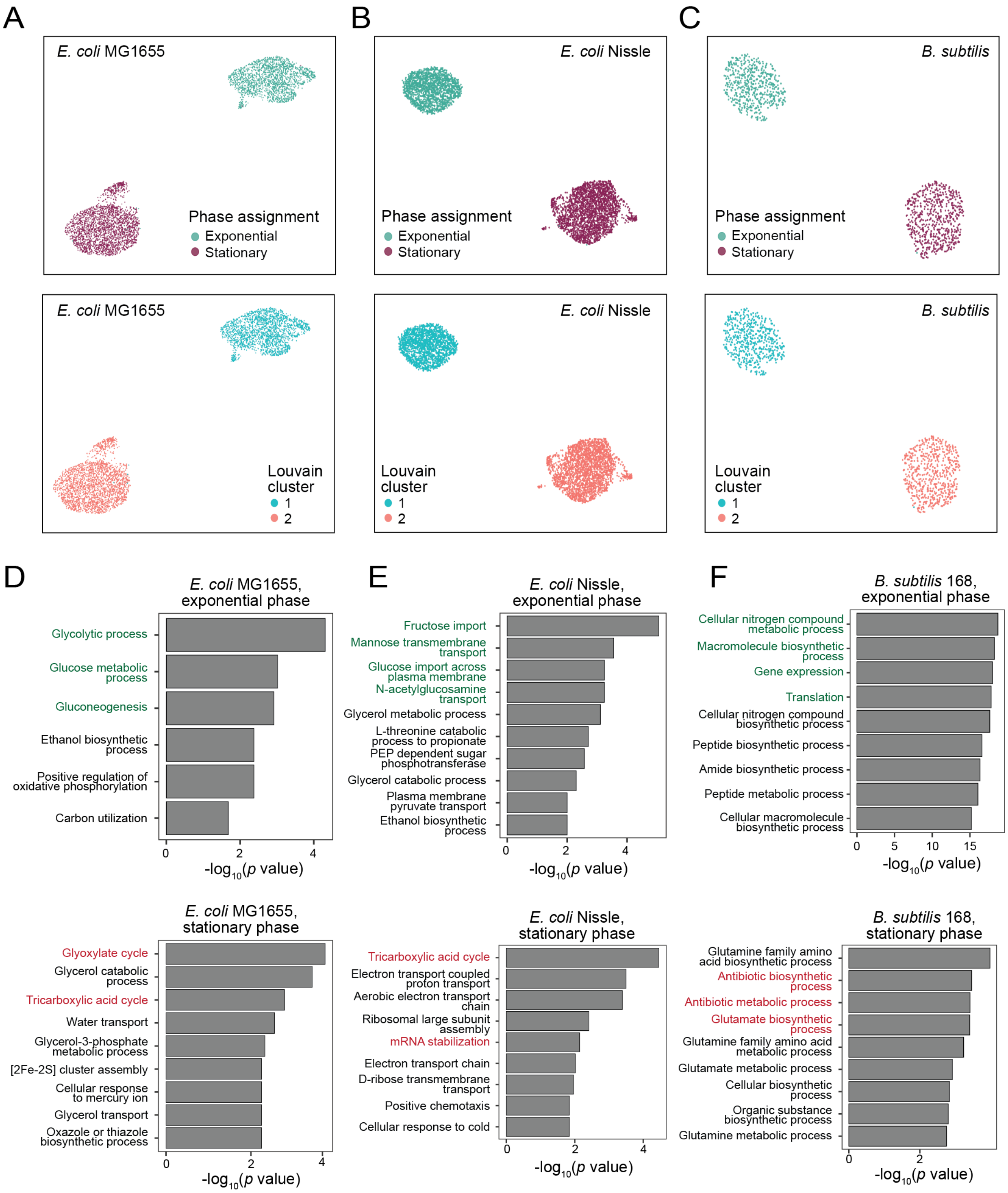
A
B
C
E. coli MG1655
Phase assignment
Exponential
Stationary
E. coli Nissle
B. subtilis
Louvain cluster
1
2
D
E. coli MG1655, exponential phase
Glycolytic process
Glucose metabolic process
Gluconeogenesis
Ethanol biosynthetic process
Positive regulation of oxidative phosphorylation
Carbon utilization
-log10(p value)
E
E. coli Nissle, exponential phase
Fructose import
Mannose transmembrane transport
Glucose import across plasma membrane
N-acetylglucosamine transport
Glycerol metabolic process
L-threonine catabolic process to propionate
PEP dependent sugar phosphotransferase
Glycerol catabolic process
Plasma membrane pyruvate transport
Ethanol biosynthetic process
F
B. subtilis 168, exponential phase
Cellular nitrogen compound metabolic process
Macromolecule biosynthetic process
Gene expression
Translation
Cellular nitrogen compound biosynthetic process
Peptide biosynthetic process
Amide biosynthetic process
Peptide metabolic process
Cellular macromolecule biosynthetic process
E. coli MG1655, stationary phase
Glyoxylate cycle
Glycerol catabolic process
Tricarboxylic acid cycle
Water transport
Glycerol-3-phosphate metabolic process
[2Fe-2S] cluster assembly
Cellular response to mercury ion
Glycerol transport
Oxazole or thiazole biosynthetic process
E. coli Nissle, stationary phase
Tricarboxylic acid cycle
Electron transport coupled proton transport
Aerobic electron transport chain
Ribosomal large subunit assembly
mRNA stabilization
Electron transport chain
D-ribose transmembrane transport
Positive chemotaxis
Cellular response to cold
B. subtilis 168, stationary phase
Glutamine family amino acid biosynthetic process
Antibiotic biosynthetic process
Antibiotic metabolic process
Glutamate biosynthetic process
Glutamine family amino acid metabolic process
Glutamate metabolic process
Cellular biosynthetic process
Organic substance biosynthetic process
Glutamine metabolic process

**Figure S4. M3-Seq profiling during exponential growth and early stationary phase. A.** UMAP projections of *E. coli* MG1655 transcriptomes in exponential and early stationary phase (top) and associated clustering (bottom, set to the lowest clustering resolution parameter). Clustering set at the lowest resolution parameter. **B.** Same as (A) but for *E. coli* Nissle. **C.** Same as (A) but for *B. subtilis* 168. **D.** GO term enrichment of select biological process calculated with marker genes identified for populations of exponential and stationary phase *E. coli* MG1655 identified in (A). Marker genes were determined by comparing the within-cluster average expression to out of cluster average expression and filtering for genes with $p$ value < 0.05 (Wilcoxon-rank sum test). The $p$-values are $-\log_{10}$ transformed such that the most strongly enriched biological processes have the highest score. Selected processes were those with the lowest $p$-values after thresholding at 0.05. Enrichments for exponential and stationary phase cells include expected processes (green and red, respectively), including growth related and energy generation processes (exponential) and those involving secondary carbon metabolism and the TCA cycle (stationary). **E.** Same as (D) but for *E. coli* Nissle. Similar to *E. coli* MG1655, enrichments include expected processes (green for exponential; red for stationary). **F.** Same as (D) but for *B. subtilis* 168. Similar to *E. coli*, enrichments include expected processes (green for exponential; red for stationary).

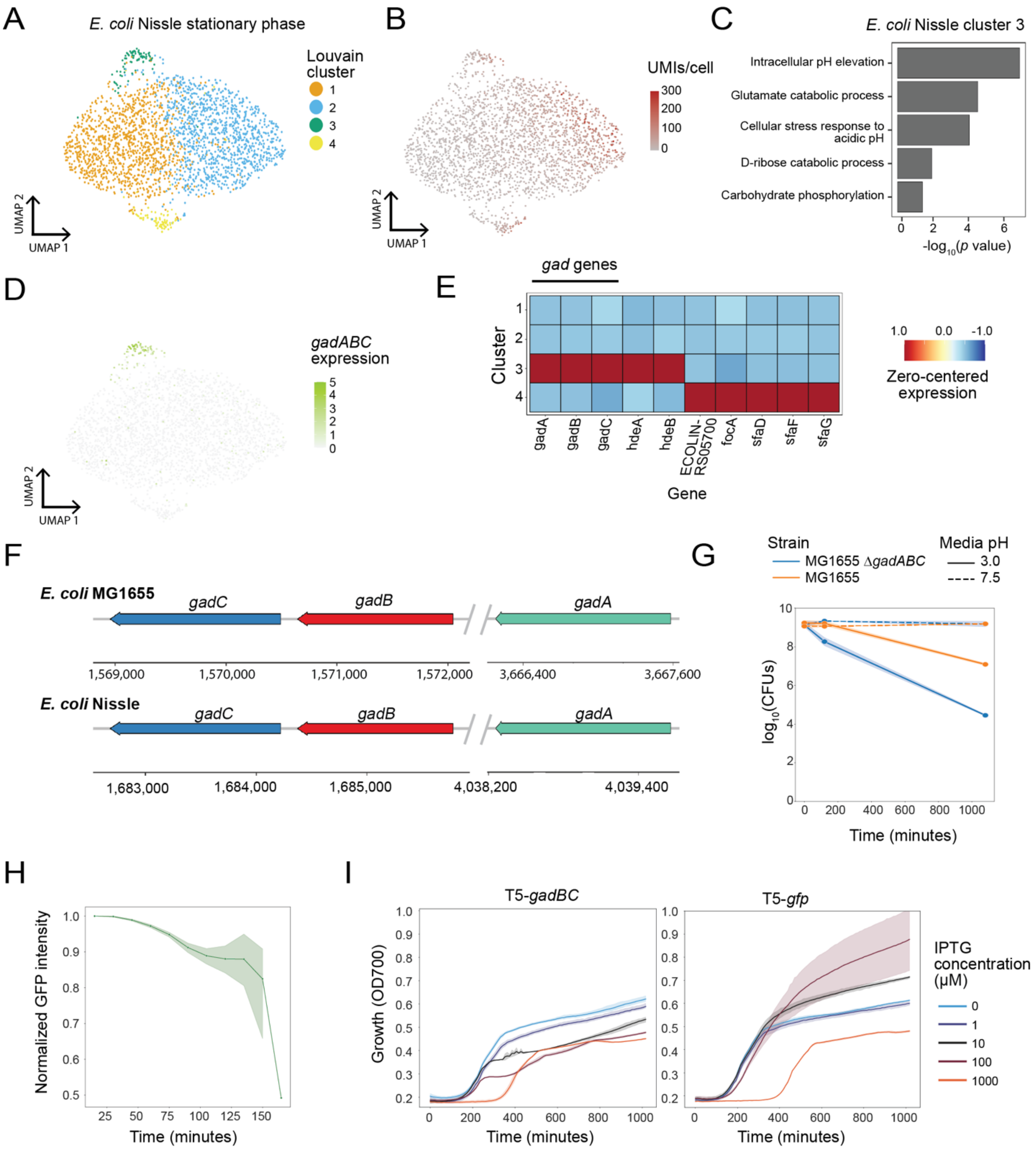
A
E. coli Nissle stationary phase
Louvain cluster
1
2
3
4
UMAP 2
UMAP 1
B
UMIs/cell
300
200
100
0
C
E. coli Nissle cluster 3
Intracellular pH elevation
Glutamate catabolic process
Cellular stress response to acidic pH
D-ribose catabolic process
Carbohydrate phosphorylation
-log10(p value)
D
gadABC expression
E
gad genes
Cluster
Gene
gadA
gadB
gadC
hdeA
hdeB
ECOLIN-RS05700
focA
sfaD
sfaF
sfaG
Zero-centered expression
F
E. coli MG1655
E. coli Nissle
gadC
gadB
gadA
1,569,000
1,570,000
1,571,000
1,572,000
3,666,400
3,667,600
1,683,000
1,684,000
1,685,000
4,038,200
4,039,400
G
Strain
MG1655 ΔgadABC
MG1655
Media pH
3.0
7.5
log10(CFUs)
Time (minutes)
H
Normalized GFP intensity
Time (minutes)
I
T5-gadBC
T5-gfp
Growth (OD700)
Time (minutes)
IPTG concentration (μM)
0
1
10
100
1000

**Figure S5. Subpopulation of early stationary phase cells expressing acid-tolerance genes also identified in *E. coli* Nissle. A.** UMAP projection of *E. coli* Nissle transcriptomes from cells at early stationary phase (OD=2.6). Colors indicate clusters of transcriptionally similar cells. **B.** Same as (A) but with color gradient indicating number of UMIs captured in each cell. **C.** GO-term enrichment of select biological processes calculated with marker genes identified for cluster 3 in (A). Marker genes were determined by comparing the within-cluster average expression to out of cluster average expression and filtering for genes with $p$ value < 0.05 (Wilcoxon-rank sum test). The $p$-values are $-\log_{10}$ transformed such that the most strongly enriched biological processes have the highest score. Selected processes were those with the lowest $p$-values (Fisher's exact test) after thresholding at 0.05. Similar to *E. coli* MG1655 (Fig. 2C), intracellular pH elevation and response to acidic pH are among the most enriched processes. **D.** Same as (A) but with color gradient indicating expression of *gadABC* genes. **E.** Zero-centered and normalized expression of marker genes for each cluster identified in (A). Marker genes were defined as those observed in at least 5% of cells in that cluster and with the lowest $p$-values (Wilcoxon rank-sum test) after thresholding to select genes with >0.5 $\log_2$ fold change between within-cluster and out-of-cluster average expression and ranking by $\log_2$ fold change. A maximum of 6 genes were included per cluster. **F.** Schematics of *gadABC* genes in the two strains of *E. coli* used in this study: MG1655 and Nissle. **G.** Plot depicts survival of wildtype *E. coli* MG1655 and Δ*gadABC* mutant with and without exposure to acid stress during early stationary phase. Compared to wildtype, Δ*gadABC* demonstrates less acid tolerance. CFUs, colony forming units. **H.** Plot depicts averaged fluorescence intensity of individual $P_{gadB}$-GFP transformed *E. coli* MG1655 cells during acid exposure normalized to the initial fluorescence of the same cell at $t$=0 ($n$ = 681 cells, error bars are 95% confidence intervals). Briefly, cells were grown to early stationary phase, transferred to an acidic pad (pH 3.0), and imaged over time. Indicating no increase in *gad* protein expression, GFP fluorescence steadily decreased during acid treatment. **I.** Plot depicts growth of *E. coli* transformed with *gadBC* (left) or GFP (right) transgene under different concentrations of IPTG inducer. While inducer levels appear unlinked to growth in GFP-expressing cells, higher amounts cause growth defects in GadBC-expressing cells.

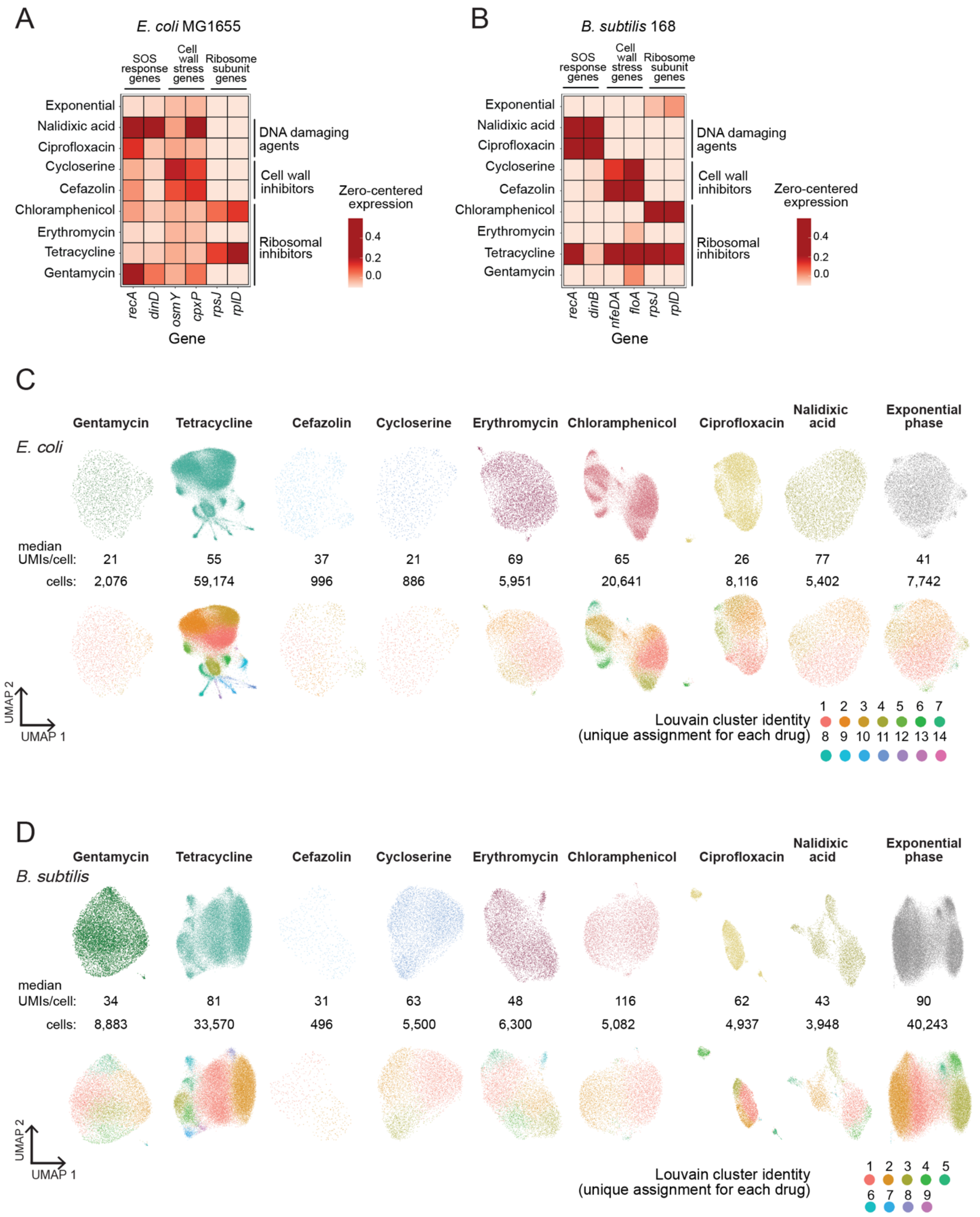
A
E. coli MG1655
SOS response genes
Cell wall stress genes
Ribosome subunit genes
Exponential
Nalidixic acid
Ciprofloxacin
Cycloserine
Cefazolin
Chloramphenicol
Erythromycin
Tetracycline
Gentamycin
DNA damaging agents
Cell wall inhibitors
Ribosomal inhibitors
Zero-centered expression
0.4
0.2
0.0
recA
dinD
osmY
cpxP
rpsJ
rplD
Gene
B
B. subtilis 168
SOS response genes
Cell wall stress genes
Ribosome subunit genes
Exponential
Nalidixic acid
Ciprofloxacin
Cycloserine
Cefazolin
Chloramphenicol
Erythromycin
Tetracycline
Gentamycin
DNA damaging agents
Cell wall inhibitors
Ribosomal inhibitors
Zero-centered expression
0.4
0.2
0.0
recA
dinB
nfeDA
floA
rpsJ
rplD
Gene
C
Gentamycin
Tetracycline
Cefazolin
Cycloserine
Erythromycin
Chloramphenicol
Ciprofloxacin
Nalidixic acid
Exponential phase
E. coli
median UMIs/cell: 21 55 37 21 69 65 26 77 41
cells: 2,076 59,174 996 886 5,951 20,641 8,116 5,402 7,742
UMAP 2
UMAP 1
Louvain cluster identity
(unique assignment for each drug)
1 2 3 4 5 6 7
8 9 10 11 12 13 14
D
Gentamycin
Tetracycline
Cefazolin
Cycloserine
Erythromycin
Chloramphenicol
Ciprofloxacin
Nalidixic acid
Exponential phase
B. subtilis
median UMIs/cell: 34 81 31 63 48 116 62 43 90
cells: 8,883 33,570 496 5,500 6,300 5,082 4,937 3,948 40,243
UMAP 2
UMAP 1
Louvain cluster identity
(unique assignment for each drug)
1 2 3 4 5
6 7 8 9

**Figure S6. Multiplexed single-cell analysis of bacterial response to eight different antibiotics. A.** Zero-centered and normalized expression of select genes in *E. coli* MG1655 cultures treated with the indicated antibiotics. Data from BW4 (Table S2). Genes were selected from among those related to the following GO terms: "Response to DNA damage", "Cell wall stress", and "Ribosome". **B.** Same as (A) but for *B. subtilis*. Genes were selected from among those related to the "Response to DNA damage" and "Ribosome" GO terms and by searching for genes known to be upregulated in response to treatment with cell wall antibiotics (i.e. cefuroxime). **C.** UMAP projections of *E. coli* MG1655 transcriptomes after treatment with indicated antibiotics (top) and corresponding cluster assignments (bottom). Clusters were uniquely defined for each population. **D.** Same as (C) but for *B. subtilis* 168.

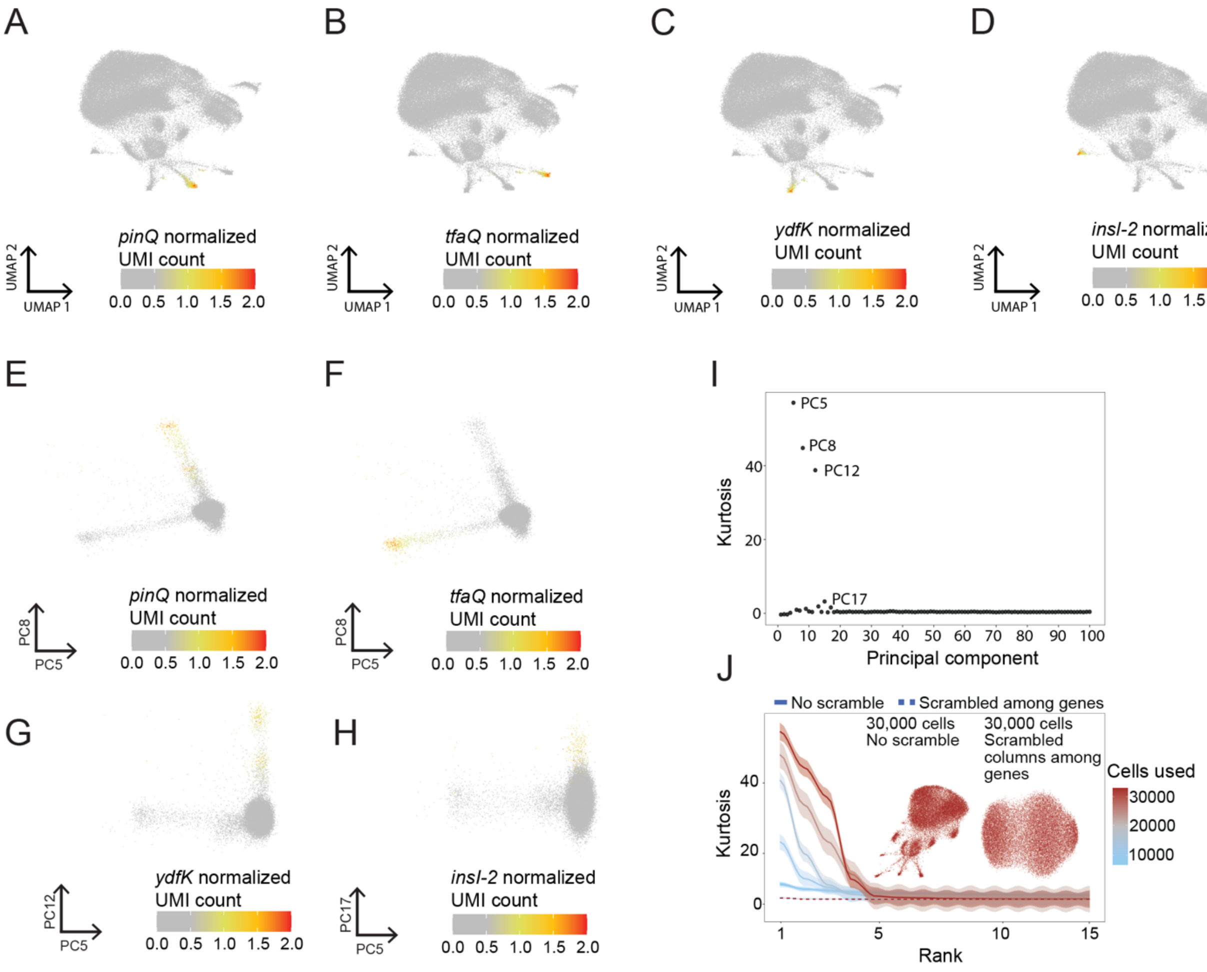
A
B
C
D
UMAP 2
UMAP 1
pinQ normalized UMI count
tfaQ normalized UMI count
ydfK normalized UMI count
insl-2 normalized UMI count
0.0 0.5 1.0 1.5 2.0
E
F
G
H
PC8
PC5
PC12
PC17
I
Kurtosis
Principal component
PC5
PC8
PC12
PC17
J
No scramble
Scrambled among genes
30,000 cells No scramble
30,000 cells Scrambled columns among genes
Cells used
30000
20000
10000
Rank

**Figure S7. Defining MGE-expressing populations of *E. coli* using M3-Seq data. A.** UMAP projection of *E. coli* MG1655 transcriptomes from cells treated with the bacteriostatic antibiotics tetracycline and chloramphenicol. Color gradient indicates normalized expression of *pinQ*, a marker gene for cluster 8 identified in Figure 3E. **B.** Same as (A) but with color gradient indicating normalized expression of *tfaQ,* a marker gene for cluster 13 identified in Figure 3E. **C.** Same as (A) but with color gradient indicating normalized expression of *ydfK*, a marker gene for cluster 12 identified in Figure 3E. **D.** Same as (A) but with color gradient indicating normalized expression of *insI-2*, a marker gene for cluster 16 identified in Figure 3E. **E.** Plots of cells in principal component space for *E. coli* treated with bacteriostatic antibiotics, wherein the color gradient indicates normalized *pinQ* expression. The principal component dimensions chosen for this analysis contained high loadings in genes that were upregulated in rare subpopulations (e.g., *pinQ*, *tfaQ*). **F.** Same as (H) but with color gradient indicating normalized *tfaQ* expression. **G.** Plots of cells in principal component space for *E. coli* treated with bacteriostatic antibiotics, wherein the color gradient indicates normalized *ydfK* expression. The principal component dimensions chosen for this analysis contained high loadings in genes enriched in rare subpopulations (e.g., *ydfK*, *tfaQ*). **H.** Same as (H) but with color gradient indicating normalized *insI-2* expression. **I.** Kurtosis of all 100 computed principal components calculated from the single cell transcriptomes of tetracycline- and chloramphenicol-treated *E. coli* MG1655. Notably, principal components with the highest kurtosis were not necessarily the same as those with the highest variance. **J.** Kurtosis of 15 principal components computed from tetracycline- and chloramphenicol-treated *E. coli* MG1655 cells, with individual lines corresponding to calculations from down-sampled subsets of cells with and without UMI counts scrambled among genes. Notably, scrambling abolishes the kurtosis signal and removes structure from clustering.

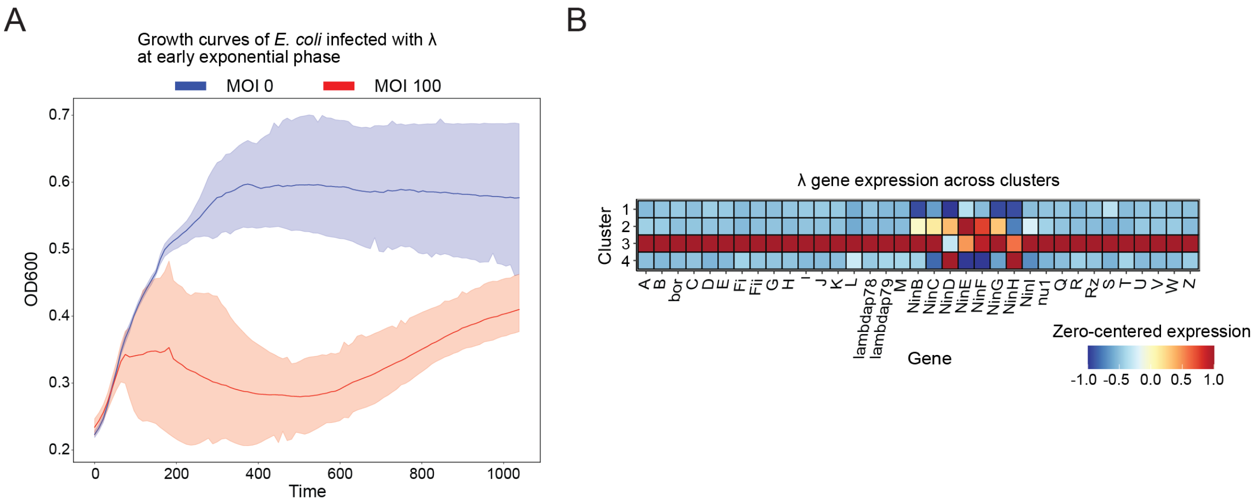


**Figure S8. Growth and gene expression in *E. coli* cells infected with λ phage. A.** Plot depicts growth of *E. coli* grown to early exponential phase (OD~0.2-0.3) and infected with λ phage (MOI ~ 100) or supplemented with phage vehicle (LB), error bars are 95% confidence intervals. **B.** Zero-centered and normalized expression of all observed λ genes for each cluster identified in Figure 5B. Genes displayed were those genes which had more than 10 UMIs across the entire population. Expression of λ genes is strongly enriched in the lytic cluster (3) but lower in the rest of the population.

| Method | Class of method | Cells profiled | Measurable genes | Median mRNA UMIs per cell | Median mRNA genes per cell | Conditions profiled |
|---|---|---|---|---|---|---|
| PETRI-Seq[8] | Plate-based | 14,975 | all expressed genes: ~4,000 per cell depending on species | 227 (*E. coli*) | 130 (*E. coli*) | 4 |
| Micro-Split[9] | Plate-Based | 25,214 | all expressed genes: ~4,000 per cell depending on species | 235 (*E. coli*)<br>397 (*B. subtilis)* | 138 (*E. coli*)<br>230 (*B. subtilis*) | 10 |
| MATQ-Seq[7] | Well-based | 126 | all expressed genes: ~4,000 per cell depending on species | >170 genes<br>(*S. typhimurium)* | 170<br>(*S. typhimurium*) | 3 |
| par-seqFISH[10] | Probe-based | 365,000 | limited by number of probes: 105 in this study | ~100<br>(*P. aeruginosa)* | 29 (*P. aeruginosa*) | 15 |
| McNulty et al.[11] | Probe-based | 3,763 | limited by number of probes: 4181 for *E. coli* & 2959 for *B. subtilis* in this study | 263 (*E. coli*)<br>325 (*B. subtilis)* | 165 (*E. coli*)<br>241 (*B. subtilis*) | 1 |

**Table S1. Statistics of existing bacterial scRNA-seq methods.** Individual run statistics from previous studies of single cell RNA-sequencing in bacteria. Numbers in each category were selected by taking maximum reported values. "Cells profiled" indicates the maximum number of bacterial cells (for any number of strains) used in a single experiment. While ultimately the limiting factor is cost per cell, all else being equal, removing rRNA reads from sequencing cost, either by using probes or rRNA depletion, will be more cost effective, with the latter also allowing for observation of unbiased gene sets. "Measurable genes" indicates the number of genes that could theoretically be measured using the indicated method. "Median mRNA UMIs per cell" indicates the maximum number of UMIs per mRNA per cells (for any number of strains and conditions) reported in the indicated study. "Median mRNA genes per cell" same as "Median mRNA UMIs per cells" except for genes. "Conditions profiled" refers to the maximum number of reported samples/conditions profiled in a single experiment. While the limiting factor in conditions profiled is time and cost, we note that indexing multiple samples using indexed primers is currently the most time and cost-efficient approach to profile multiple conditions.

| Experiment | Cells loaded | Single cell transcriptomes recovered | Species | Library treatment |
|---|---|---|---|---|
| BW1 | 100,000 | 13,416 | *E. coli* (MG1655), *B. subtilis* (168) | No rRNA depletion |
| BW2 | 37,500 | 4,975 | *E. coli* (MG1655), *B. subtilis* (168) | *Post hoc* rRNA depletion |
| BW3 | 67,670 | 15,538 | *E. coli* (MG1655), *E. coli* (Nissle), *B. subtilis* (168) | *Post hoc* rRNA depletion |
| BW4 | 426,000; 585,000 | 107,310; 122,361 | *E. coli* (MG1655), *B. subtilis* (168) | *Post hoc* rRNA depletion |

**Table S2. Details of the single-cell sequencing experiments presented in this study.** Statistics of the single-cell sequencing experiments presented in this study. "Cells loaded" refers to the number of single cells loaded into one lane of the Chromium Controller. "Single cell transcriptomes recovered" were the number of single-cell transcriptomes that passed the UMI threshold (*Materials and Methods*). "Species" refers to the species analyzed in each experiment. BW4 also included *P. aeruginosa* PA14 and *S. aureus* MRSA but cell / UMI recovery was low and reads mapping to these species were discarded. "Library treatment" refers to additional steps performed during library preparation.

**Table S3. Sample statistics from all experiments included in study.** BW4 also included *P. aeruginosa* PA14 and *S. aureus* MRSA but cell / UMI recovery was low and reads mapping to these species were discarded

**Table S4. Oligonucleotides used in this study.** Due to the large number of oligonucleotides used in this study, each workbook is split into the different purposes that the oligonucleotide was used for, which are “RT oligos”, “rRNA depletion oligos”, and “General oligos”.

**Table S5. Marker genes for all the clusters and samples analyzed in this study.** Each workbook here refers to marker genes for a single species (*E. coli* MG1655, *E. coli* Nissle, *B. subtilis* 168) within all the conditions profiled within that single-cell sequencing experiment (BW1, BW2, BW3, BW4).

**Movie S1. Movie of *E. coli* MG1655 recovering from the acid-stress recovery assay.** Representative movie of the acid-stress recovery assay (Fig. 2G) conducted with *E. coli* MG1655 transformed with $P_{gadB}$-GFP. Briefly, cells were treated with acid (pH 3.0) in early stationary phase for one hour and then transferred to a fresh LB pad for imaging. GFP and phase channels are overlaid.

**Movie S2. Movie of *E. coli* MG1655 during strong acid treatment.**
Representative movie of *E. coli* MG1655 transformed with $P_{gadB}$-GFP during acid treatment (pH 3.0). Briefly, cells were grown to early stationary phase, transferred to an acidic pad (pH 3.0), and imaged over time. Indicating no increase in *gad* protein expression, GFP fluorescence steadily decreased during acid treatment. GFP and phase channels are overlaid.
